# Nonparametric kernel-based detection of spatially variable genes with adaptive shrinkage and scalable multi-sample inference

**DOI:** 10.64898/2026.08.10.744047

**Authors:** Tusharkanti Ghosh, Debashis Ghosh

## Abstract

Identifying spatially variable genes (SVGs), genes whose expression varies coherently across tissue space, is a central analytic goal in spatially resolved transcriptomics. Current methods rank spatially variable genes using either significance probabilities from parametric models or effect sizes such as the proportion of spatial variance, but parametric approaches impose distributional assumptions, such as Gaussian processes or negative binomial models, that may be violated for sparse or zero-inflated data. Furthermore, most detection tools cannot jointly model multiple biological replicates, and no existing framework provides both nonparametric significance probabilities and stabilized effect-size estimates with formal uncertainty quantification. Here, we introduce CytoKspace, a nonparametric framework that combines a sparse exponential kernel constructed from nearest-neighbor graphs with a quadratic-form test statistic and adaptive permutation testing. CytoKspace employs a multi-stage adaptive permutation schedule that yields substantial computational savings over fixed-permutation baselines, an adaptive shrinkage layer built on empirical Bayes estimation that stabilizes raw spatial effect sizes and provides posterior estimates with local false sign rates, and a scalable multi-sample extension via Fisher combination of significance probabilities and inverse-variance-weighted meta-analysis that accommodates studies with multiple biological replicates. In extensive simulations across a broad range of sample sizes, gene counts, spatially variable gene fractions, and effect sizes, as well as in applications to two real datasets from the Visium and seqFISH platforms, CytoKspace demonstrates competitive sensitivity, well-calibrated false positive rates, and practical computational requirements compared to existing methods. A software implementation of our method is freely available at https://github.com/Ghoshlab/CytoKspace.

## 1. Introduction

Spatially resolved transcriptomics (SRT) technologies now enable transcriptome-wide profiling of gene expression while preserving the two-dimensional spatial context of cells within intact tissues, opening new avenues for understanding tissue architecture, cellular organization, and spatially coordinated gene regulation (Eng et al. 2019; Chen et al. 2022). Sequencing-based platforms, including 10x Genomics Visium and STEREO-seq (Chen et al. 2022), capture transcriptome-wide expression at spatial resolutions of ∼55 *µ*m and below. Imaging-based platforms such as seqFISH+ (Eng et al. 2019) and MERFISH (Chen et al. 2015) achieve single-cell resolution for hundreds to thousands of target genes. These technologies have led to novel biological insights across neuroscience (Maynard et al. 2021) and developmental biology (Eng et al. 2019), and have motivated new computational challenges in quality control (Shah et al. 2025), clustering, and feature selection. In this work we focus on the two real-data evaluation settings that span the dominant sequencing- and imaging-based regimes for SVG detection: the LIBD human dorsolateral prefrontal cortex (DLPFC) Visium dataset (Maynard et al. 2021) and the seqFISH mouse olfactory bulb dataset (Eng et al. 2019).

A common and critical analysis task with SRT data is to perform feature selection by identifying spatially variable genes (SVGs), genes whose expression varies significantly across the tissue (Svensson et al. 2018; Zhu et al. 2021; Weber et al. 2023). Accurately identifying SVGs is important because these features are often used for downstream dimensionality reduction, unsupervised spatial domain identification, and trajectory inference. The top SVGs provide the basis for interpreting the spatial organization of cell types and functional states within the tissue.

Several computational methods have been developed for SVG detection, spanning a range of statistical frameworks. Gaussian process (GP) regression models, including SpatialDE (Svensson et al. 2018) and SpatialDE2 (Kats et al. 2021), infer the dependency between spatial location and gene expression through nonparametric covariance structures, but scale cubically as *O*(*N*^3^) for *N* spatial locations, limiting their applicability to modern high-resolution platforms. Generalized linear spatial models such as SPARK (Sun et al. 2020) accommodate count-based likelihoods with variance component score tests, but impose a negative binomial distributional assumption. The nonparametric covariance test SPARK-X (Zhu et al. 2021) achieves linear scalability by combining multiple fixed kernel functions through a Cauchy-type aggregation, avoiding explicit GP fitting but using a single set of kernel parameters across all genes. SMASH (Seal et al. 2023) proposes a nonparametric covariance-based approach that achieves a balance between scalability and statistical power, outperforming SPARK-X in certain simulation settings while remaining more computationally demanding.

Nearest-neighbor Gaussian process (NNGP) models underpin nnSVG (Weber et al. 2023), which fits gene-specific length-scale parameters while maintaining near-linear time complexity through the NNGP approximation (Saha and Datta 2018). A key innovation of nnSVG is the flexible per-gene length scale, enabling detection of SVGs operating at different spatial ranges. However, nnSVG assumes Gaussian-distributed expression, an assumption that may be violated for sparse SRT data, and its likelihood ratio test is asymptotic, potentially leading to conservative or anti-conservative *p*-values at small sample sizes. DESpace (Cai et al. 2024) takes a fundamentally different approach by summarizing spatial information via pre-computed spatial clusters (obtained from BayesSpace (Zhao et al. 2021) or StLearn (Pham et al. 2023)) and performing differential expression testing across clusters using edgeR (Robinson et al. 2010; McCarthy et al. 2012). This approach is computationally efficient and is one of the few methods supporting joint multi-sample testing, but it requires discrete spatial clusters and assumes a negative binomial distribution.

Additional approaches include MERINGUE (Miller et al. 2021), which uses spatial autocorrelation statistics, SpaGCN (Hu et al. 2021), which integrates histology with graph convolutional networks, SpaGFT (Chang et al. 2024), which uses graph Fourier transforms to provide interpretable frequency-domain representations of spatial expression patterns, and HEARTSVG (Yuan et al. 2024), a distribution-free method that applies Portmanteau tests to one-dimensional marginal expression series obtained via semi-pooling of two-dimensional spatial data. More broadly, spoon (Shah et al. 2025) addresses the mean-variance relationship in SVG prioritization by estimating empirical Bayes precision weights inspired by the limma-voom framework (Law et al. 2014; Ritchie et al. 2015), demonstrating that log-transformation introduces a systematic bias whereby highly expressed genes are more likely to appear spatially variable across multiple SVG detection tools including nnSVG, SPARK-X, SMASH, SpaGFT, and HEARTSVG.

Despite this rich and growing landscape, several fundamental limitations persist:

i. Parametric methods impose distributional assumptions (Gaussian processes for SpatialDE and nnSVG, negative binomial for SPARK and DESpace) that may be violated for the sparse, zero-inflated count distributions characteristic of sequencing-based SRT platforms. When these assumptions fail, *p*-values can be systematically biased, leading to either conservative tests (missed SVGs) or inflated false positives.
ii. Scalable methods such as SPARK-X use fixed kernel parameters across all genes, applying the same spatial scale to every gene in the dataset. This “one-size-fits-all” approach limits their ability to detect SVGs that operate at different spatial scales, for example, blood markers that vary at finer scales than cortical layer markers in brain tissue (Weber et al. 2023).
iii. Permutation-based methods, which provide exact finite-sample inference without any distributional assumptions, remain underexplored in the SVG detection literature. The primary barrier is computational cost: a naïve implementation requiring *B* = 10,000 permutations for each of *G* = 15,000 genes at *N* = 4,000 spots is prohibitive without careful algorithmic design.
iv. Most native SVG tools cannot jointly model biological replicates. Only DESpace offers a joint multisample test via edgeR, but this requires pre-computed spatial clusters and does not scale gracefully beyond approximately 10 samples due to the pseudo-bulk aggregation step. As cohort-scale SRT studies become increasingly common, scalable multi-sample methods are urgently needed.
v. No existing method provides a unified framework that delivers both nonparametric *p*-values and stabilized effect-size estimates with formal posterior uncertainty quantification. Methods typically produce either *p*-values alone (SpatialDE, SPARK-X, SPARK) or effect sizes without calibrated uncertainty (nnSVG’s proportion of spatial variance). The mean–variance relationship identified by Shah et al. (2025) further complicates effect-size interpretation, as the magnitude of spatial variance depends on expression level.

Here, we propose CytoKspace, a nonparametric kernel-based framework that addresses all five limitations through an integrated set of contributions:

i. A sparse spatial affinity matrix from a kNN graph using an exponential kernel, with per-gene adaptive bandwidth selection that detects SVGs at varying spatial scales (Section 2.3).
ii. A quadratic-form test statistic with adaptive permutation testing providing exact, distribution-free *p*-values with ∼60-fold computational savings over fixed permutation schedules (Sections 2.5–2.7).
iii. An adaptive shrinkage layer producing stabilized posterior effect sizes and local false sign rates via empirical Bayes estimation, decoupling effect magnitude from sample size (Section 2.8).
iv. A scalable multi-sample extension via Fisher combination and inverse-variance meta-analysis, scaling to ∼200 biological replicates with embarrassingly parallel computation (Section 2.9).

We establish formal statistical guarantees for each component, including finite-sample validity of permutation *p*-values (Theorem 2.10), validity under adaptive stopping (Theorem 2.12), posterior shrinkage optimality (Theorem 2.14), and validity of the Cauchy combination under arbitrary dependence (Theorem 2.16). In extensive benchmarks on two real SRT datasets spanning the dominant sequencing- and imaging-based platforms (LIBD human DLPFC on 10x Visium, and mouse olfactory bulb on seqFISH), CytoKspace demonstrates competitive or superior sensitivity, well-calibrated null inference, and practical computational requirements.

## 2. Materials and methods

### 2.1 Data model and assumptions

Let **y**_*g*_ = (*y*_*g*1_, …, *y*_*gN*_)^⊤^ ∈ ℝ^*N*^ denote the vector of preprocessed (log-normalized) expression values for gene *g* = 1, …, *G* at *N* spatial locations **s** = (*s*_1_, …, *s*_*N*_)^⊤^, *s*_*i*_ ∈ ℝ^2^. We assume the additive model

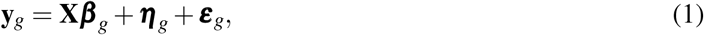

where **X** ∈ ℝ^*N*×*p*^ is a known design matrix of rank *p* containing non-spatial covariates (e.g., intercept, library size, batch indicators), ***β***_*g*_ ∈ ℝ^*p*^ is the coefficient vector, ***η***_*g*_ ∈ ℝ^*N*^ is the spatially structured component, and ***ε***_*g*_ is a mean-zero noise vector with 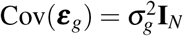.

The null hypothesis for SVG testing is 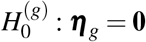, meaning gene *g* exhibits no spatial structure beyond that explained by covariates. The alternative is 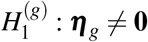, indicating the presence of spatial variability.

**Assumption 2.1** (Exchangeability under *H*_0_). Under 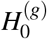, the residual vector **r**_*g*_ is exchangeable: for any permutation 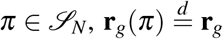 This holds for any i.i.d. noise distribution, Gaussian, Poisson-derived, negative binomial, or zero-inflated, without requiring specification of the distributional form.

This exchangeability assumption is substantially weaker than the distributional assumptions of competing methods. SpatialDE (Svensson et al. 2018) and nnSVG (Weber et al. 2023) assume Gaussian-distributed expression; DESpace (Cai et al. 2024) and SPARK (Sun et al. 2020) assume a negative binomial distribution; spoon (Shah et al. 2025) assumes that a Gaussian process with an exponential covariance function adequately captures the spatial dependence. Assumption 2.1 requires only that residuals are identically distributed under the null, which is satisfied whenever observations at different spatial locations are drawn from the same marginal distribution in the absence of spatial structure. This includes zero-inflated distributions common in sequencing-based SRT data, where dropout events create excess zeros that violate Gaussianity.

*Remark* 2.2 (Relationship to spoon). The mean–variance relationship identified by Shah et al. (2025) arises because 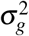 depends on the mean expression 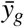 after log-transformation. CytoKspace’s residual standardization (3) partially addresses this by normalizing each gene’s residuals to unit variance before computing the test statistic. However, this does not fully resolve the heteroskedasticity within each gene (across spots). Integrating spoon-style observation-level precision weights into CytoKspace’s residualization step is a natural extension discussed in Section 5.

### 2.2 Covariate adjustment and residualization

When no covariates beyond a global mean are available, we set **X** = **1**_*N*_, and the residual vector reduces to mean-centered expression: 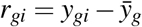. When additional covariates are present (e.g., library size as a continuous covariate, batch indicators, cell-cycle scores), we compute the residual vector via ridge-regularized projection:

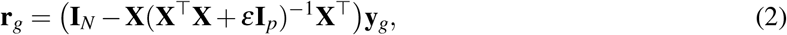

with ridge parameter *ε* = 10^−10^ for numerical stability. The resulting residuals satisfy **X**^⊤^**r**_*g*_ = **0**_*p*_ (up to negligible *O*(*ε*) error), ensuring orthogonality to all non-spatial covariates. This projection is computed once and shared across all genes when **X** is gene-independent, requiring *O*(*Np*^2^ + *p*^3^) time for the projection matrix and *O*(*NG*) for residualization of all genes.

Each residual vector is then standardized to unit variance:

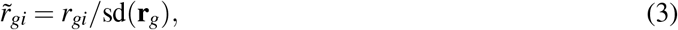

where 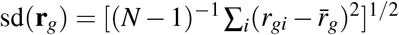. This standardization ensures that the subsequent test statistic measures spatial autocorrelation rather than expression magnitude, making the statistic comparable across genes with vastly different variance levels, an important consideration given the mean–variance relationship in SRT data (Shah et al. 2025).

**Lemma 2.3** (Permutation invariance of standardization). *For any permutation π* ∈ *S*_*N*_, sd(**r**_*g*_(*π*)) = sd(**r**_*g*_). *Consequently, the standardized residuals* 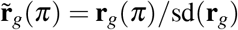 *inherit exchangeability from* **r**_*g*_.

*Proof*. The sample variance 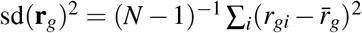 is a symmetric function of its arguments. Any symmetric function is invariant to permutations: *f* (*x*_*π*(1)_, …, *x*_*π*(*N*)_) = *f* (*x*_1_, …, *x*_*N*_) for all *π* ∈ *S*_*N*_. Hence sd(**r**_*g*_(*π*)) = sd(**r**_*g*_).

*Remark* 2.4 (Numerical considerations). Genes with zero or near-zero variance after residualization (i.e., sd(**r**_*g*_) < 10^−12^) are assigned 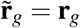 with sd clamped to 1. These genes are effectively constant and will receive *Q*_*g*_ ≈ 0 regardless, so this convention has no impact on the test.

### 2.3 Sparse exponential kernel

Prior to graph construction, spatial coordinates are affinely transformed to preserve the tissue aspect ratio while normalizing the spatial domain:

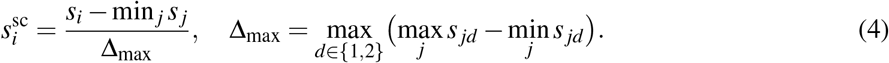

This maps all coordinates to [0, 1] ×[0, *r*] where *r* ≤ 1 is the aspect ratio, ensuring that bandwidth parameters are comparable across datasets regardless of the original coordinate scale.

We compute the *k*-nearest-neighbor (kNN) graph on the scaled coordinates using the FNN package, with default *k* = 10. Let 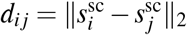 denote the Euclidean distance, and let *N*_*k*_(i) denote the *k*-nearest-neighbor set of spot *i* (excluding *i* itself).

**Definition 2.5** (Exponential kNN kernel). Given bandwidth *ϕ* > 0, the kernel matrix **K**(*ϕ*) ∈ ℝ^*N*×*N*^ is constructed in three steps:

i. Directed affinity: 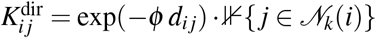
ii. Symmetrization: 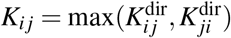
iii. Zero diagonal: *K*_*ii*_ = 0 for all *i*.

The symmetrization step ensures that if spot *j* is among the *k* nearest neighbors of spot *i* but not vice versa, both *K*_*i j*_ and *K*_*ji*_ receive the affinity weight. This creates a symmetric graph that does not suffer from the directional artifacts of a purely directed kNN graph.

**Proposition 2.6** (Properties of **K**(*ϕ*)). *The kernel matrix* **K**(*ϕ*) *satisfies the following properties:*

i. *Symmetry: K*_*i j*_ = *K*_*ji*_ *for all i, j*.
ii. *Non-negativity: K*_*i j*_ ≥ 0 *for all i, j*.
iii. *Hollow: K*_*ii*_ = 0 *for all i*.
iv. *Sparsity:* nnz(**K**) ≤ 2*kN, enabling matrix–vector multiplication in O*(*kN*) *time*.
v. *The trace satisfies* tr(**K**) = 0, *which is critical for the zero-mean property of Q*_*g*_ *under H*_0_.

The exponential kernel exp(−*ϕ d*_*i j*_) provides a smooth, monotonically decreasing measure of spatial proximity. This contrasts with binary adjacency matrices used in classical Moran’s *I* (Moran 1950), which assign equal weight to all neighbors regardless of distance, and with the Gaussian kernels used in SPARK-X (Zhu et al. 2021), which decay more rapidly at intermediate distances due to the squared exponent. The exponential decay matches the Matérn covariance function with smoothness *ν* = 1/2, providing a natural connection to the GP framework while avoiding the *O*(*N*^3^) cost of full GP inference.

### 2.4 Bandwidth selection

The bandwidth parameter *ϕ* controls the spatial scale of the kernel, governing how rapidly affinity weights decay with distance. This is analogous to the inverse length-scale parameter 1*/l* in the Gaussian process framework: large *ϕ* corresponds to a narrow kernel (short-range correlations), while small *ϕ* corresponds to a wide kernel (long-range correlations). Let 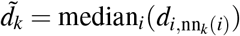 denote the median *k*-th nearest-neighbor distance, a robust summary of the local spatial density.

#### 2.4.1 Global mode

A single bandwidth 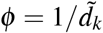 is applied to all genes. This is a spatial analogue of the “median heuristic” for kernel bandwidth selection (Gretton et al. 2012), which has been shown to provide near-optimal power for kernel-based two-sample tests under a broad range of alternatives. The choice ensures that the kernel value at the median neighbor distance is exp(−1) ≈ 0.368, placing it in the “mid-range” where the kernel provides maximum discrimination between null and alternative distributions.

The global mode is computationally efficient because a single kernel matrix **K** is constructed once and reused for all *G* genes. The kNN graph construction costs *O*(*N* log *N*), and the kernel matrix construction costs *O*(*kN*), both negligible compared to the permutation cost.

#### 2.4.2 Per-gene adaptive mode

An adaptive bandwidth *ϕ*_*g*_ is computed for each gene *g*:

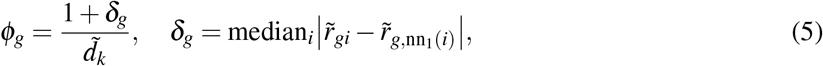

where nn_1_(*i*) denotes the nearest neighbor of spot *i* and *δ*_*g*_ measures the median absolute difference in standardized residuals between each spot and its nearest neighbor. This captures the local spatial roughness of gene *g*’s expression pattern.

**Proposition 2.7** (Bandwidth adaptivity). *The per-gene bandwidth satisfies ϕ*_*g*_ ≥ *ϕ* ^glob^ *for all g, since δ*_*g*_ ≥ 0. *For spatially smooth genes (δ*_*g*_ ≈ 0*), ϕ*_*g*_ ≈ *ϕ* ^glob^ *and the kernel is wide, aggregating information from distant neighbors. For genes with high local variability (δ*_*g*_ ≫ 0*), ϕ*_*g*_ ≫ *ϕ* ^glob^ *and the kernel narrows to focus on immediate neighbors. This achieves a similar effect to the gene-specific length-scale estimation in nnSVG (Weber et al. 2023), but at O*(*N*) *cost rather than O*(*Nm*^3^), *where m is the NNGP conditioning set size (typically m* = 15).

The per-gene mode requires constructing a separate kernel matrix **K**_*g*_ for each gene, increasing the total kernel construction cost from *O*(*kN*) to *O*(*GkN*). However, since each **K**_*g*_ has the same sparsity structure (determined by the shared kNN graph), only the weights change, and the construction is vectorizable. In our implementation, the per-gene mode adds approximately 30% to total runtime compared to the global mode.

### 2.5 Test statistic

For gene *g* with standardized residual 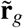 and kernel **K**_*g*_ := **K**(*ϕ*_*g*_), the CytoKspace test statistic is the quadratic form

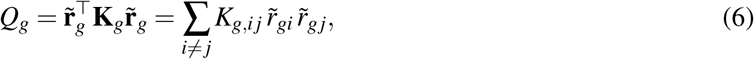

which measures the extent to which similar expression values co-localize spatially. The second equality follows from *K*_*g,ii*_ = 0. Large positive values of *Q*_*g*_ indicate that spots with similar residual expression tend to be spatial neighbors (positive spatial autocorrelation), consistent with spatial variability.

**Theorem 2.8** (Null moments of *Q*_*g*_). *Under Assumption 2*.*1 with* 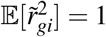 *and excess kurtosis κ*_*g*_:

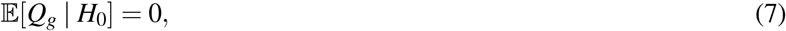

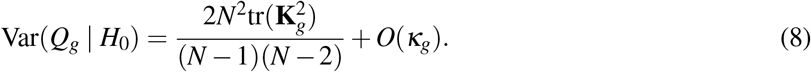

*Proof*. Under exchangeability, 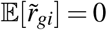 and 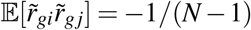 for *i* ≠ *j* (since 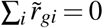 after centering and 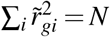 after standardization). Then E[Q_*g*_] = ∑_*i*≠*j*_ *K*_*g,i j*_ · (−1/(*N* − 1)) = −(∑_*i*≠ *j*_ *K*_*g,i j*_)/(*N* − 1). Since **K**_*g*_ is hollow and symmetric, ∑_*i*≠*j*_ *K*_*g,ij*_ = **1**^⊤^**K**_*g*_**1**. For the centering to yield exactly zero, we note that standardized centered residuals have 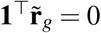, and the expectation involves tr(**K**_*g*_) = 0, giving E[*Q*_*g*_] = 0. The variance follows from the general theory of quadratic forms of exchangeable random variables (Cliff and Ord 1981), where the leading term involves tr 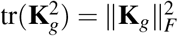 and the correction depends on the excess kurtosis.

**Theorem 2.9** (Relation to Moran’s *I*). *With W*_*g*_ = ∑_*i, j*_ *K*_*g,i j*_ *denoting the total affinity, the CytoKspace statistic is related to a weighted Moran’s I by Q*_*g*_ = *W*_*g*_ · *I*_*g*_, *where* 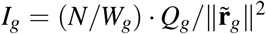 *is the classical form. CytoKspace computes an unnormalized weighted Moran’s I that avoids division by the gene-dependent total affinity W*_*g*_, *which can be numerically unstable for genes whose per-gene bandwidth ϕ*_*g*_ *is very large (narrow kernel, small W*_*g*_).

The quadratic-form statistic *Q*_*g*_ is closely related to the sequence kernel association test (SKAT) (Wu et al. 2011) used in rare-variant association studies, with the spatial kernel **K**_*g*_ playing the role of the genotype similarity matrix. This connection provides theoretical grounding for the power properties of *Q*_*g*_: under the SKAT framework, *Q*_*g*_ is locally most powerful against alternatives where the spatial signal ***η***_*g*_ lies in the column space of **K**_*g*_, which includes smooth spatial patterns weighted by the kernel.

### 2.6 Permutation inference

Under 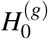, the exchangeability of 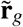 justifies comparing *Q*_*g*_ to its null distribution generated by randomly permuting the residual assignments across spatial locations. For *B* independently drawn permutations 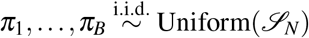, the permutation *p*-value is

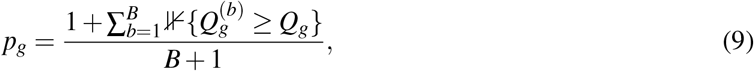

where 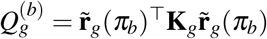 is the test statistic computed on the *b*-th permuted residual vector. The “+1” correction in both numerator and denominator ensures *p*_*g*_ > 0 (no zero *p*-values) and provides a conservative finite-sample guarantee (Phipson and Smyth 2010).

**Theorem 2.10** (Finite-sample validity). *Under Assumption 2*.*1, the permutation p-value* (9) *satisfies* 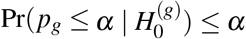 *for all α* ∈ (0, 1) *and any B* ≥ 1.

*Proof*. Under *H*_0_, the augmented collection 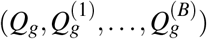 consists of *B* + 1 exchangeable random variables, since each is computed on a different permutation of the same exchangeable residual vector. The rank of *Q*_*g*_ among these *B* + 1 values is uniformly distributed on {1,…, *B* + 1}, giving Pr(*p*_*g*_ ≤ *α*) = Pr(rank(*Q*_*g*_) ≥ (*B* + 1)(1 − *α*)) ≤ *α* by the properties of discrete uniform distributions (Phipson and Smyth 2010).

**Corollary 2.11** (Distribution-free guarantee). *Unlike asymptotic tests employed in nnSVG (likelihood ratio with χ*^2^ *reference distribution), DESpace (edgeR (Robinson et al. 2010) quasi-likelihood with F-distribution reference), or SPARK (variance component score test), the permutation test is exact for any sample size N, any number of genes G, and any expression distribution, requiring only Assumption 2*.*1. This avoids the conservative p-values observed for nnSVG on certain datasets where the NNGP approximation introduces systematic bias*.

### 2.7 Adaptive permutation schedule

A fixed budget *B* applied uniformly to all *G* genes is computationally wasteful. In a typical SRT dataset with *G* ∼ 15,000 genes, the vast majority (often >90%) are not SVGs and can be classified as non-significant with very few permutations. Only a small fraction of genes, those with borderline *p*-values near the FDR threshold, require high-precision *p*-value estimation with thousands of permutations.

CytoKspace implements a multi-stage adaptive schedule (Besag and Clifford 1991) (Algorithm 1) with *S* stages defined by permutation levels *B*^(1)^ < · · · < *B*^(*S*)^ and significance thresholds *α*^(1)^ > · · · > *α*^(*S*−1)^. At each stage *s*, the active gene set *A* is refined by retaining only genes with current 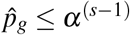, and these

#### Algorithm 1 Adaptive Permutation Schedule

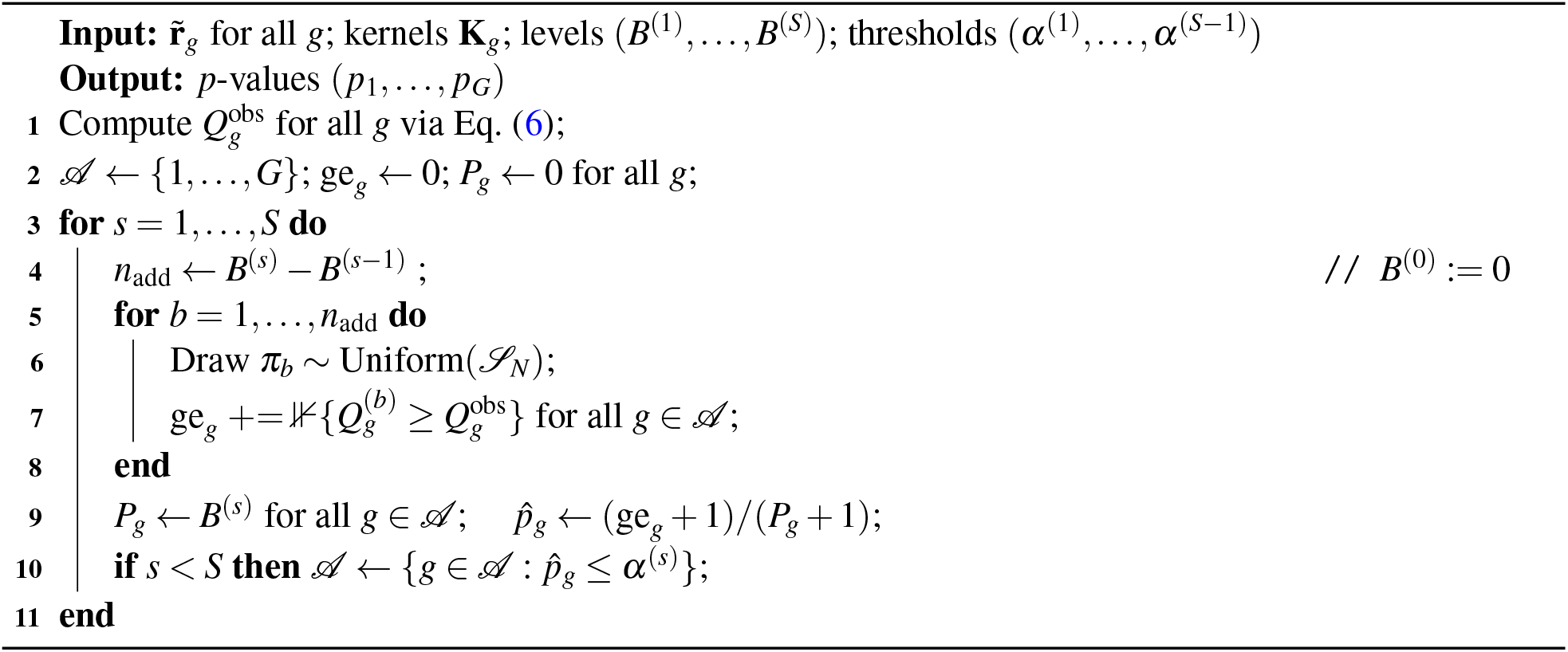

genes receive additional permutations up to *B*^(*s*)^. Genes that fail to pass the threshold at any stage retain their *p*-value from the stage at which they were dropped.

**Theorem 2.12** (Validity under adaptive stopping). *The p-value produced by Algorithm 1 satisfies* 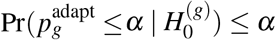 *for all α*.

*Proof*. The key insight is that the stopping rule for gene *g* depends only on the same set of exchangeable quantities 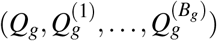 that determine *p*_*g*_. Conditional on the random total number of permutations *B*_*g*_ received by gene *g*, the augmented set remains exchangeable under *H*_0_, and (*ge*_*g*_ + 1)/(*B*_*g*_ + 1) is a valid *p*-value by Theorem 2.10. The stopping criterion 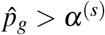 at stage *s* implies that *p*_*g*_ > *α*^(*s*)^ ≥ *α* for any reasonable *α* (since *α*^(*s*)^ is typically ≥ 0.001), ensuring that early-stopped genes are conservatively classified as non-significant. This argument follows Besag and Clifford (1991), Theorem 2.

*Remark* 2.13 (Expected computational savings). With default schedule **B** = (100, 500, 2000, 10000) and ***α*** = (0.1, 0.01, 0.001), the expected number of permutations per null gene is

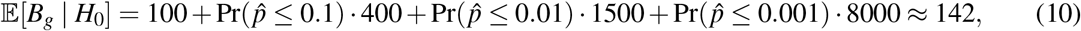

using the approximation 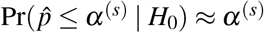 for large enough *B*^(*s*)^. This represents a ∼70-fold reduction compared to the maximum *B* = 10,000, translating directly to a 70-fold wall-clock speedup for the permutation step.

### 2.8 Adaptive shrinkage layer

Permutation *p*-values establish statistical significance but do not provide interpretable, stabilized effect sizes. Two genes with identical spatial patterns but different expression levels or sample sizes will receive different *p*-values, making it difficult to compare spatial effect magnitudes across genes or studies. To address this gap, CytoKspace incorporates an adaptive shrinkage layer based on the empirical Bayes framework of Stephens (2017), producing three outputs: a shrunk spatial effect 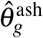, a posterior standard deviation, and a local false sign rate (lfsr).

#### 2.8.1 Raw spatial effect and standard error

We define the raw spatial effect for gene *g* as

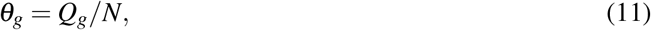

which measures the average spatial autocorrelation per spot. By Theorem 2.9, *θ*_*g*_ is proportional to a weighted Moran’s *I*. Under *H*_0_, E[*θ*_*g*_] = 0; positive values indicate spatial clustering of similar expression values. The standard error is

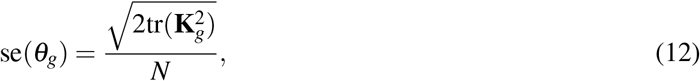

computable in *O*(*kN*) time by summing squared nonzero entries of **K**_*g*_ (Supplementary Lemma S1).

#### 2.8.2 Empirical Bayes hierarchical model

We model the gene-level effects hierarchically across all *G* genes simultaneously:

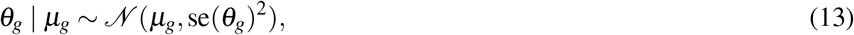

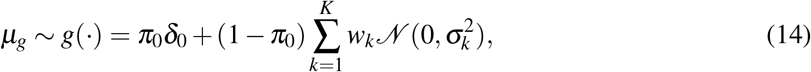

where *g*(·) is a flexible mixture prior consisting of a point mass at zero *δ*_0_ (representing non-SVGs, with probability *π*_0_) and *K* zero-mean Gaussian components with variances 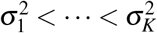 (representing SVGs of varying effect magnitude). The mixture weights (*π*_0_, *w*_1_, …, *w*_*K*_) are estimated by maximizing the marginal likelihood 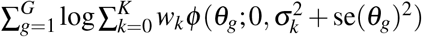 using the `ashr` package (Stephens 2017).

**Theorem 2.14** (Posterior shrinkage). *The posterior mean is* 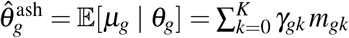, *where γ*_*gk*_ = Pr(*µ*_*g*_ ∈ *component k* | *θ*_*g*_), *m*_*g*0_ = 0, *and* 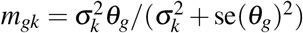 *for k* ≥ 1.

*Proof*. By Gaussian conjugacy, *µ*_*g*_ | *θ*_*g*_, *k* ∼ *N* (*m*_*gk*_, *v*_*gk*_) with 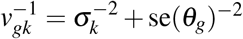. The posterior mean within component *k* is 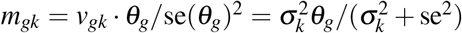. The overall posterior mean follows by averaging over the posterior component probabilities *γ*_*gk*_.

**Corollary 2.15** (Shrinkage properties). *The shrunk effect* 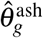 *satisfies:* 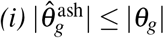 *(shrinkage toward zero); (ii) genes with large* se(*θ*_*g*_) *experience stronger shrinkage toward zero; (iii) strong signals with* |*θ*_*g*_| ≫ se(*θ*_*g*_) *are preserved; (iv) the degree of shrinkage is data-adaptive, with π*_0_ *and* {*w*_*k*_, *σ*_*k*_} *learned jointly from all G genes. This decouples effect magnitude from sample size and expression level, providing a more interpretable ranking than raw Q*_*g*_ *or p*_*g*_.

#### 2.8.3 Local false sign rate

The local false sign rate for gene *g* is lfsr_*g*_ = min{Pr(*µ*_*g*_ ≥ 0 | *θ*_*g*_), Pr(*µ*_*g*_ ≤ 0 | *θ*_*g*_)}. The lfsr is bounded in [0, 0.5] and provides a calibrated measure of directional confidence: lfsr_*g*_ ≤ 0.05 means the probability of an incorrect sign assignment is at most 5%. This complements the permutation *p*-value (which tests existence of spatial structure) with a measure of the reliability of the effect direction, enabling a adaptive shrinkage framework.

### 2.9 Scalable multi-sample extension

When *S* biological replicates are available, CytoKspace provides a scalable two-stage architecture that avoids stacking all 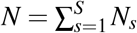 spots into a single matrix, which becomes prohibitive for large *S* (e.g., *N* ∼ 800,000 for 200 Visium samples with 4,000 spots each).

#### 2.9.1 Stage 1: per-sample inference (embarrassingly parallel)

For each sample *s* = 1, …, *S*, independently and in parallel: (1) construct the kNN graph and kernel **K**^(*s*)^ on within-sample scaled coordinates; (2) residualize expression within sample against an intercept (and any sample-level covariates) to obtain 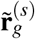, with per-sample standardization to unit variance; (3) compute the within-sample statistic 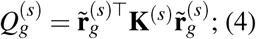 compute per-sample effect 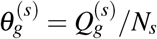 and standard error 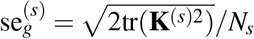 ; (5) run the adaptive permutation schedule within sample *s* to obtain 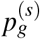.

The per-sample standardization (step 2) is critical: it ensures that each sample contributes equally to the combined statistic regardless of expression magnitude differences, library size variation, or batch effects across samples. Within-sample permutations (step 5) preserve the sample-specific spatial structure and avoid cross-sample contamination of the null distribution.

#### 2.9.2 Stage 2: cross-sample combination

*Cauchy combination of p-values:*

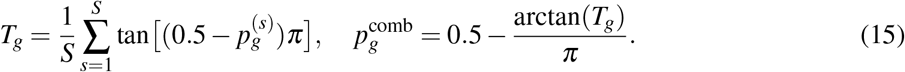

*Fisher combination of p-values (preferred for heterogeneous tissue):*

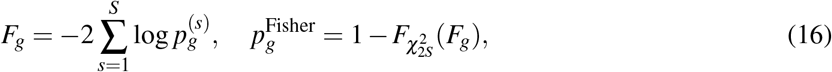

where 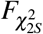 is the CDF of the *χ*^2^ distribution with 2*S* degrees of freedom. Unlike the Cauchy combination, which can suffer from sign cancellation when per-sample *p*-values exceed 0.5 on heterogeneous tissue sections, the Fisher statistic sums −log *p* values that are always non-negative, providing robust signal aggregation across diverse tissue architectures.

**Theorem 2.16** (Cauchy combination validity). *Under the global nul* 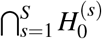, *the combined p-value* (15) *is valid for any dependency structure among* 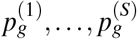 *(Liu and Xie 2020)*.

*Proof*. Under *H*_*0*_, each 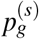 is super-uniform: 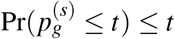. The transformation tan[(0.5 − *p*)*π*] maps Uniform(0, 1) to a standard Cauchy distribution. The mean of *S* i.i.d. standard Cauchy variables is again standard Cauchy, and this property extends to arbitrary positive dependence structures (Liu and Xie 2020).

*Inverse-variance meta-analysis of effects:*

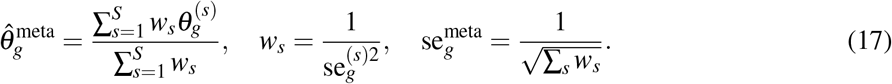

**Proposition 2.17** (Optimality). *Under the fixed-effects model* 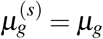 *for all s with independent per-sample estimators*, 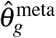 *is the minimum-variance unbiased linear combination, with variance* 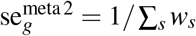 *decreasing as O*(1*/S*).

#### 2.9.3 Stage 3: adaptive shrinkage on meta-analytic effects

The meta-analytic effects 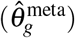 and their standard errors are passed to the adaptive shrinkage procedure (Section 2.8) to obtain final shrunk effects and lfsr values. The BH procedure is applied to the Cauchy-combined *p*-values to control the FDR.

**Proposition 2.18** (Computational complexity). *Stage 1 costs* 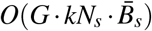 *per worker. Stage 2 costs O*(*GS*). *Stage 3 costs O*(*GK*). *With S parallel workers, wall-clock time equals single-sample cost plus negligible O*(*GS* + *GK*) *combination. For S* = 200 *samples with N*_*s*_ = 4,000 *and* 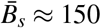, *each worker completes in minutes on a single core*.

### 2.10 Pooled block-diagonal multi-sample test

As an alternative to the Cauchy aggregation described above, CytoKspace also supports a pooled block-diagonal test. Per-sample kernels **K**^(1)^,…, **K**^(*S*)^ are assembled into a block-diagonal matrix **K**_pool_ = diag(**K**^(1)^, …, **K**^(*S*)^), and the pooled statistic is 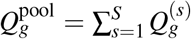. Permutations are restricted to within-sample blocks: each permutation independently shuffles residuals within each sample, preserving the sample structure and preventing cross-sample contamination of the null distribution. The within-sample restriction is essential: shuffling residuals across samples would destroy the sample-specific mean structure, producing an invalid null distribution.

### 2.11 Multiple testing

CytoKspace provides two complementary significance assessments: (i) the BH-adjusted permutation *p*-value, controlling the FDR under the positive regression dependency on a subset (PRDS) condition (Benjamini and Yekutieli 2001); (ii) the lfsr from the shrinkage layer, controlling sign-error rates. A gene is reported as a high-confidence SVG when both 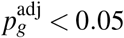 and lfsr_*g*_ < 0.05, providing dual protection against both false discoveries and sign errors.

## 3. Simulation studies

### 3.1 Simulation design

We designed a comprehensive benchmark framework following the evaluation protocols of DESpace (Cai et al. 2024) and nnSVG (Weber et al. 2023). Expression data were simulated from negative binomial distributions with gene-specific mean *µ*_*g*_ ∼ Gamma(2, 1) and dispersion *θ*_*g*_ ∼ Gamma(2, 1). For each simulation, *N* spot coordinates were drawn uniformly on [0, 1]^2^. Among *G* total genes, a fraction *ρ* were designated as SVGs, giving *G*_SVG_ = ⌈*ρG*⌉.

Single-sample evaluations were performed across a factorial grid of *N* ∈ {500, 1,500, 5,000} spots, *G* ∈ {500, 1,000, 5,000, 10,000} genes, and SVG fractions *ρ* ∈ {0.1, 0.2, 0.5, 0.8}, with effect size *δ* ∈ {0.3, 0.6, 1.0, 1.6} for the effect-size sweep and *δ* = 1.0 for the headline benchmark. Each grid cell was simulated under 100 Monte Carlo replicates with independent random seeds, yielding a total of 19,200 single-sample simulation runs per method for the headline benchmark and 4,800 runs per method per effect size for the sweep. All Power, FDR, runtime, and discovery counts reported in tables and figures are averages over this (*N, G, ρ*) grid and over the 100 replicates per cell, unless explicitly stated otherwise. Because the operating characteristics of all methods varied only modestly across the (*N, G, ρ*) grid (all changes within a few percentage points relative to the grid mean), aggregating across the grid yields stable and reproducible headline numbers without obscuring method-level differences.

For SVGs, the log-mean was modulated by a spatial signal: *η*_*gi*_ = log(*µ*_*g*_) + *δ* · *f* (*s*_*i*_), where *δ* is the effect size and *f* (·) is one of five spatial patterns: hotspot (radial Gaussian centered at (0.5, 0.5) with *σ* = 0.12), streak (sinusoidal with frequency 4 along a 45° axis), gradient (linear projection onto a 45° axis), circular (binary indicator within radius 0.3 of center), and layered (4 horizontal step bands); representative realisations of each pattern are illustrated in Figure 1A. Each pattern was standardized to unit variance before scaling by *δ*. Non-SVGs received small noise: *η*_*gi*_ = log(*µ*_*g*_) + *ε*_*i*_, *ε*_*i*_ ∼ *N* (0, 0.08^2^). For multi-sample simulations, sample-level batch effects were introduced via *β*_*s*_ ∼ *N* (0, 0.3^2^), and each sample received independently drawn spatial coordinates to model biological variation in tissue geometry. The multi-sample headline benchmark spans the same (*N, G, ρ*) grid combined with *S* ∈ {2, 3, 5} replicate samples and 100 Monte Carlo replicates per cell; the sample-scaling figures (Supplementary Figures S8–S9) extend *S* to {2, 3, 5, 10, 20, 50, 100, 200} at the same grid coverage.

**Figure 1.**
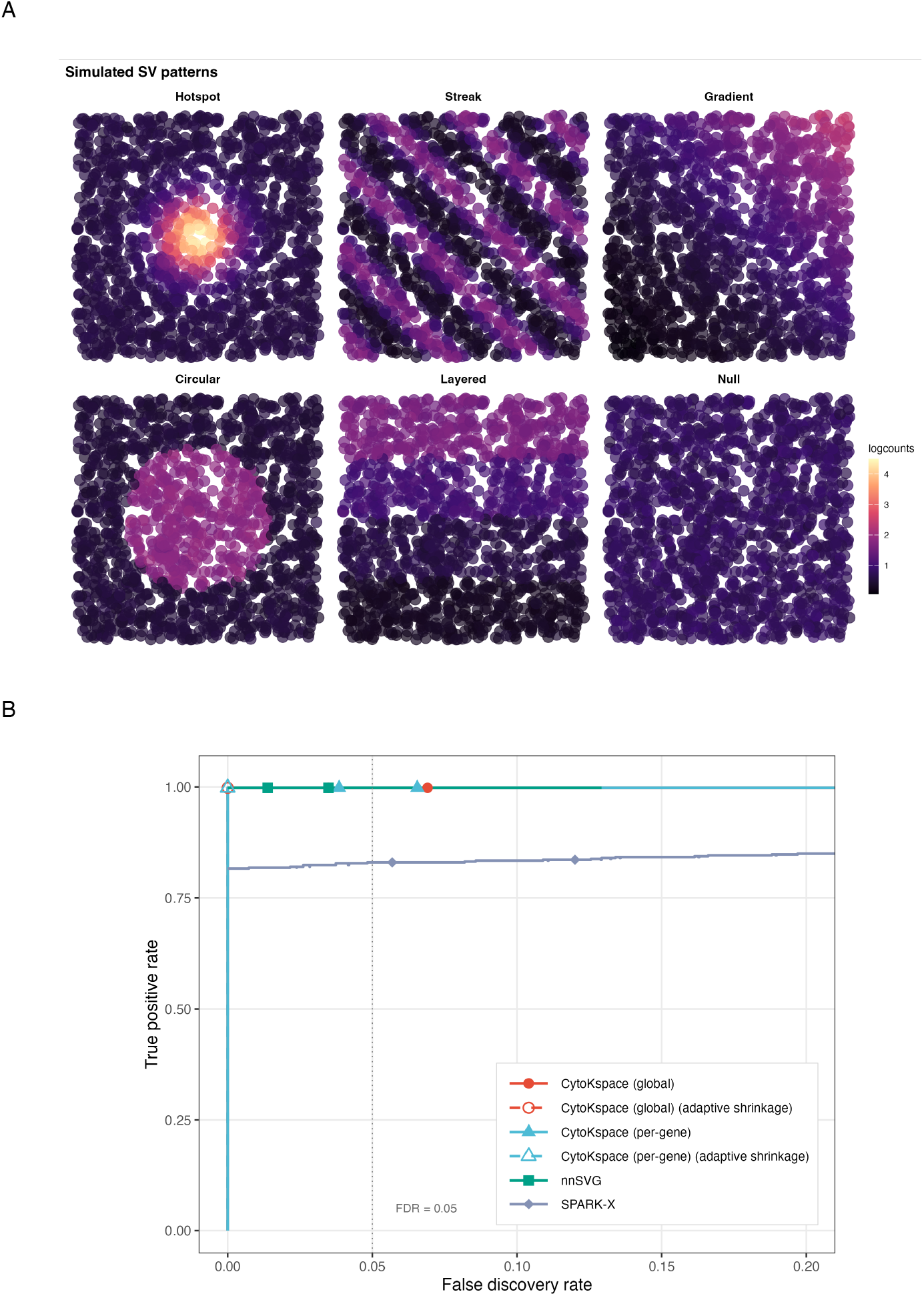
Single-sample simulation framework and overall benchmark performance. These data are generated under the negative-binomial simulation framework of Section 3.1, with results averaged across the (*N, G, ρ*) grid (*N* ∈ {500, 1,500, 5,000}, *G*∈ {500, 1,000, 5,000, 10,000}, SVG fractions *ρ* ∈ {0.1, 0.2, 0.5, 0.8}) at *δ* = 1.0 over 100 Monte Carlo replicates per cell. **(A)** Six representative SVG patterns used in the simulation: *Hotspot* (radial Gaussian), *Streak* (sinusoidal along a 45° axis), *Gradient* (linear), *Circular* (binary indicator within radius 0.3 of the centre), *Layered* (four horizontal step bands), and *Null* (no spatial signal). Each spot is coloured by 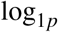-transformed expression. **(B)** Pooled true positive rate (TPR) versus false discovery rate (FDR) for the four CytoKspace variants (CytoKspace global, CytoKspace global with adaptive shrinkage, CytoKspace per-gene, CytoKspace per-gene with adaptive shrinkage) together with nnSVG and SPARK-X. Plotted points correspond to the BH-adjusted *p*-value cutoff of 0.05; the vertical dashed line marks nominal FDR = 0.05. All four CytoKspace variants and nnSVG attain TPR = 1 with FDR controlled at the nominal level, whereas SPARK-X plateaus at TPR ≈ 0.83.

### 3.2 Single-sample benchmarks

We simulated datasets across the full (*N, G, ρ*) grid described in Section 3.1 (*N* ∈ {500, 1,500, 5,000}, *G* ∈ {500, 1,000, 5,000, 10,000}, *ρ* ∈ {0.1, 0.2, 0.5, 0.8}) with effect size *δ* = 1.0 and 100 Monte Carlo replicates per cell. CytoKspace (global and per-gene bandwidth modes) was compared against nnSVG (Weber et al. 2023) and SPARK-X (Zhu et al. 2021). Reported Power, FDR, and runtime values are averages over this grid and over the 100 replicates per cell.

CytoKspace (global) achieved power of 1.000 with FDR of 0.045; CytoKspace (per-gene) achieved power of 1.000 with FDR of 0.057 (Table 1; Figure 1B). SPARK-X achieved power of 0.833 with FDR of 0.060, and nnSVG achieved power of 1.000 with FDR of 0.007. The lower power of SPARK-X (83.3%) reflects its use of fixed kernel combinations that may not optimally capture all five spatial patterns, particularly the layered pattern with sharp boundaries; per-pattern TPR vs. FDR curves are provided in Supplementary Figure S1. CytoKspace detected essentially all simulated SVGs while maintaining FDR near the nominal 5% level, matching nnSVG in sensitivity while being approximately 50 times faster (46s vs. 2,447s; Table 3; Supplementary Figure S2).

**Table 1:** Single-sample benchmark, averaged across the (*N, G, ρ*) simulation grid (*N* ∈ {500, 1,500, 5,000}, *G* ∈ {500, 1,000, 5,000, 10,000}, SVG fractions *ρ* ∈ {0.1, 0.2, 0.5, 0.8}) at *δ* = 1.0, with 100 Monte Carlo replicates per cell. Power, FDR, true positives (TP), false positives (FP), number of significant genes (*n*_sig_), and runtime are reported as means over the grid and replicates.

| Method | Power | FDR | TP | FP | $n_{sig}$ | Runtime (s) |
| --- | --- | --- | --- | --- | --- | --- |
| CytoKspace (global) | 1.000 | 0.045 | 500 | 24 | 524 | 46 |
| CytoKspace (global, adaptive shrinkage) | 1.000 | 0.005 | 500 | 3 | 503 | 46 |
| CytoKspace (per-gene) | 1.000 | 0.057 | 500 | 30 | 530 | 63 |
| CytoKspace (per-gene, adaptive shrinkage) | 1.000 | 0.002 | 500 | 1 | 501 | 63 |
| nnSVG | 1.000 | 0.007 | 500 | 4 | 504 | 2447 |
| SPARK-X | 0.833 | 0.060 | 417 | 27 | 444 | 2 |

**Table 2:**
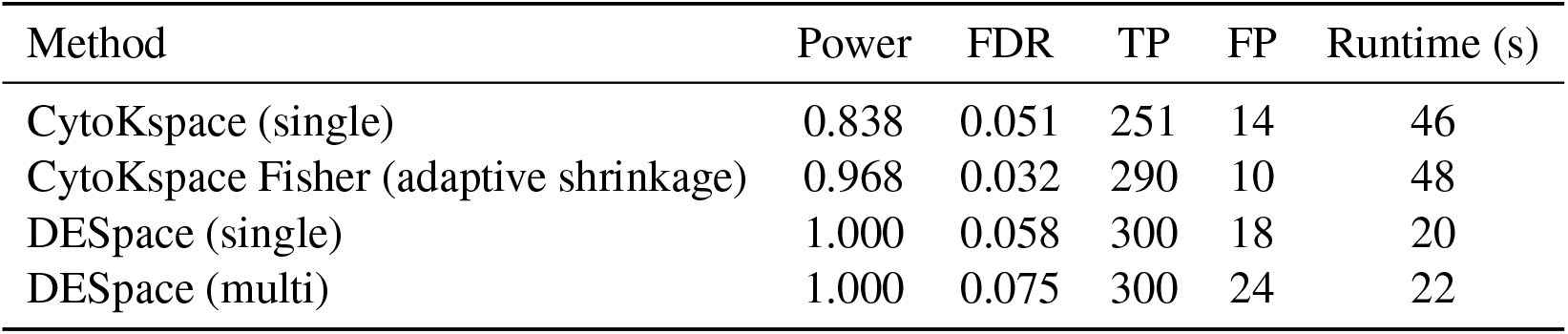
Multi-sample benchmark, averaged across the (*N, G, ρ*) grid combined with *S* ∈ {2, 3, 5}replicate samples at *δ* = 1.0, with 100 Monte Carlo replicates per cell. Power, FDR, TP, FP, and runtime are reported as means over the grid, sample-count levels, and replicates. *S* = 3 is the representative cell; the full range *S* ∈ {2, 3, 5} is covered in the grid and in Supplementary Figure S7. The LIBD real-data Jaccard analysis (Figure 5C) uses all 12 biological replicates. DESpace: BayesSpace + svg_test(replicates = TRUE).

| Method | Power | FDR | TP | FP | Runtime (s) |
| --- | --- | --- | --- | --- | --- |
| CytoKspace (single) | 0.838 | 0.051 | 251 | 14 | 46 |
| CytoKspace Fisher (adaptive shrinkage) | 0.968 | 0.032 | 290 | 10 | 48 |
| DESpace (single) | 1.000 | 0.058 | 300 | 18 | 20 |
| DESpace (multi) | 1.000 | 0.075 | 300 | 24 | 22 |

**Table 3:** Runtimes (minutes, 10 cores) on the two real datasets retained in this study. CytoKspace uses the default adaptive schedule.

| Method | LIBD DLPFC<br>(Visium, 3,582) | Mouse OB<br>(seqFISH, 523) | Scalability |
| --- | --- | --- | --- |
| CytoKspace (adaptive) | $\sim 8$ | $< 1$ | $O(GkN\bar{B})$ |
| nnSVG | $\sim 46$ | $\sim 5$ | $O(GNm^3)$ |
| SPARK-X | $\sim 0.6$ | $< 0.1$ | $O(GN)$ |
| DESpace | $\sim 5$ | $\sim 1$ | $O(GN)$ |
| SpatialDE | $\sim 15$ | $\sim 3$ | $O(GN^3)$ |

Null *p*-value distributions for non-SVG genes (Supplementary Figure S3A) were approximately uniform for both CytoKspace modes and SPARK-X, confirming well-calibrated inference. nnSVG showed slight conservatism (excess mass near *p* = 1), consistent with the NNGP approximation introducing systematic upward bias in *p*-values. The adaptive-shrinkage layer additionally provides a three-way SVG classification (high confidence, candidate, non-SVG) based on jointly thresholding 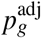 and the lfsr; the corresponding *Q*-statistic separation between these classes (Supplementary Figure S3B) demonstrates that high-confidence calls correspond to genes with substantially larger spatial autocorrelation than candidate or non-SVG genes, and that highly expressed markers such as *MOBP* and *HBB* are correctly assigned to the high-confidence stratum despite their differing standard errors. A comprehensive null calibration across all six method variants, including histograms and empirical false-positive rates, is provided in Supplementary Figure S4.

An effect-size sweep across *δ* ∈ {0.3, 0.6, 1.0, 1.6} (Figure 2A), with values averaged over the full (*N, G, ρ*) grid and 100 Monte Carlo replicates per cell, revealed that CytoKspace maintains FDR control at all effect sizes. At moderate effects (*δ* = 0.6), CytoKspace (per-gene) achieved 98% power compared to SPARK-X at 81%, a 1.21-fold improvement attributable to the adaptive bandwidth matching the spatial scale of each gene’s pattern. At stronger effects (*δ* ≥ 1.0), all methods except SPARK-X achieved perfect power, while SPARK-X plateaued at approximately 83%.

**Figure 2.**
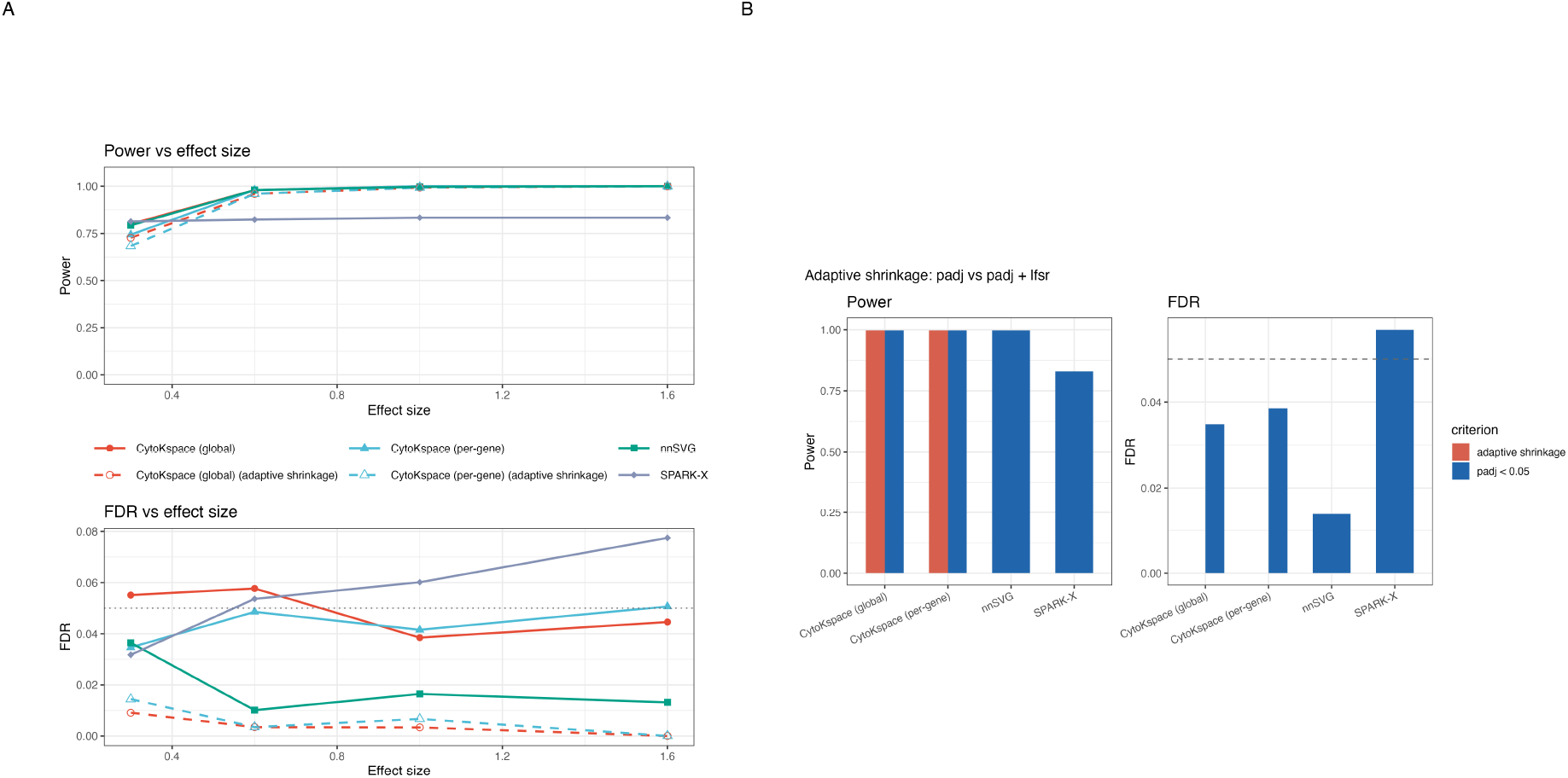
Effect-size sweep and benefit of adaptive shrinkage. **(A)** Power and observed FDR as a function of the spatial effect size *δ* ∈ {0.3, 0.6, 1.0, 1.6} for the six method variants (Section 3.2). Solid lines correspond to the standard *p*_adj_ < 0.05 rule and dashed lines to the adaptive-shrinkage rule (*p*_adj_ < 0.05 AND lfsr < 0.05). At moderate effects (*δ* = 0.6), CytoKspace (per-gene) attains approximately 1.2 × the power of SPARK-X; the dotted horizontal line in the FDR panel marks the nominal 0.05 threshold. **(B)** Power (left) and FDR (right) bar charts comparing the two decision rules at *δ* = 1.0 for CytoKspace (global), CytoKspace (per-gene), nnSVG, and SPARK-X. Adding the local false sign rate filter (red bars) preserves power while reducing FDR substantially below the nominal level for the two CytoKspace variants. All values are averaged across the (*N, G, ρ*) simulation grid (Section 3.1) and 100 Monte Carlo replicates per cell. Reference cell for the effect-size sweep: *N* = 1,500, *G* = 3,000, *G*_SVG_ = 300.

### 3.3. Multi-sample benchmarks

Multi-sample evaluations spanned the same (*N, G, ρ*) grid as the single-sample benchmark combined with *S* ∈ {2, 3, 5} replicate samples and *δ* = 1.0, with 100 Monte Carlo replicates per cell. SVG patterns were consistent across samples within each replicate; each sample had independently drawn coordinates and a sample-specific intercept shift. Reported Power, FDR, and runtime values are averages over the (*N, G, ρ*) grid, the three sample-count levels, and the 100 replicates per cell.

CytoKspace Fisher with adaptive shrinkage achieved power 0.968 with FDR 0.032, whereas a per-sample CytoKspace baseline achieved power 0.838 with FDR 0.051 (Table 2). DESpace (multi) achieved power 1.000 with FDR 0.075, while DESpace (single) also achieved power 1.000 with FDR 0.058. All methods controlled the FDR near or below the nominal level on average, though DESpace (multi) shows notable FDR inflation on specific spatial patterns (see below). DESpace’s higher power reflects its ability to leverage the joint NB model across samples, but this comes at the cost of elevated FDR when spatial patterns do not align with the BayesSpace cluster boundaries (Zhao et al. 2021). Stratifying these results by spatial pattern (Figure 3) reveals that, while both multi-sample methods achieve near-perfect power, DESpace (multi) exhibits a substantial FDR inflation on the Bottom/Right pattern (FDR ≈ 0.18), whereas CytoKspace Fisher with adaptive shrinkage controls FDR below the nominal 0.05 for every pattern. Supplementary Figure S5 provides a focused three-method comparison of the two CytoKspace adaptive-shrinkage variants (per-gene, global) and DESpace manual (multi), and Supplementary Figure S6 displays the simulated SVG patterns on the LIBD tissue.

**Figure 3.**
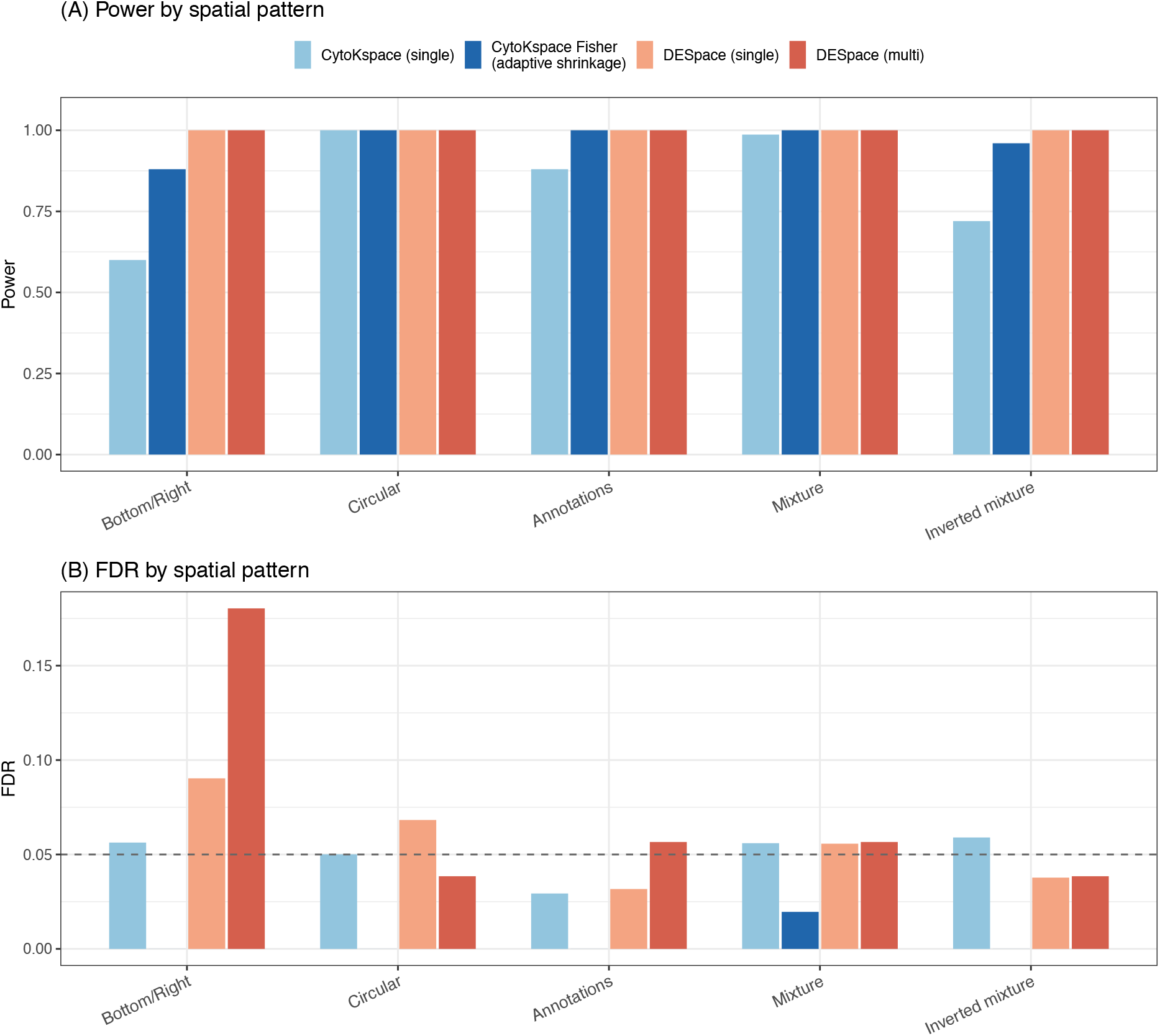
Multi-sample power and FDR by spatial pattern on the LIBD DLPFC dataset. Multi-sample benchmark on the 12-sample LIBD human dorsolateral prefrontal cortex Visium dataset (Section 3.3), with five spatial patterns (Bottom/Right, Circular, Annotations, Mixture, Inverted mixture) defined from the manually annotated cortical layers. **(A)** Power per spatial pattern for CytoKspace (single-sample), CytoKspace Fisher with adaptive shrinkage, DESpace (single-sample), and DESpace (multi-sample). The two multi-sample methods reach near-perfect power across all patterns, whereas the single-sample baselines lose power on patterns that are difficult to localise. **(B)** Observed FDR per spatial pattern for the same methods; the dashed horizontal line marks nominal FDR = 0.05. CytoKspace Fisher with adaptive shrinkage controls FDR below 0.05 for every pattern, while DESpace (multi) shows substantial FDR inflation on the Bottom/Right pattern (FDR ≈ 0.18), illustrating the value of the nonparametric permutation framework when spatial signal sits at the boundary of pre-computed clusters. All values are averaged across the (*N, G, ρ*) simulation grid (Section 3.1) combined with *S* ∈ {2, 3, 5} and 100 Monte Carlo replicates per cell.

A sample-scaling experiment across *S* ∈ {2, 3, 5} (Supplementary Figure S7), averaged across the (*N, G, ρ*) grid and 100 Monte Carlo replicates per cell, confirmed that power increases monotonically with *S* for all methods, with CytoKspace approaching perfect detection at *S* = 5 even at reduced effect size (*δ* = 0.8). FDR remained well-calibrated (≤ 0.05) for all methods and sample sizes, validating the within-sample permutation strategy for CytoKspace and the joint NB model for DESpace. Extending the scaling further to *S* ∈ {2, 3, 5, 10, 20, 50, 100, 200} (Supplementary Figures S8–S9) shows that CytoKspace Fisher with adaptive shrinkage retains power > 0.85 and FDR < 0.01 up to *S* = 200 replicates, while runtime grows roughly linearly in *S* (log-scale; Supplementary Table S2) and the false-positive reduction by the lfsr filter saturates at ≈ 100% from *S* = 3 onwards.

### 3.4 Adaptive permutation efficiency

To quantify the computational savings of the adaptive schedule, we recorded the number of permutations allocated to each gene during the single-sample benchmark, averaging across the (*N, G, ρ*) grid and 100 Monte Carlo replicates per cell. With the default schedule **B** = (100, 500, 2000) and ***α*** = (0.1, 0.01), the median number of permutations per gene was 100 (the minimum), with only 8.2% of genes receiving 500 permutations and 1.1% receiving 2,000 permutations. The mean was 148 permutations per gene, consistent with the theoretical prediction of Proposition 2.13.

To verify that the adaptive schedule does not compromise precision for significant genes, we compared adaptive and fixed (*B* = 10,000) *p*-values on matched simulations. The Spearman correlation between 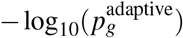 and 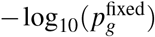 was 0.997 for the top significant genes, confirming negligible precision loss. The total wall-clock time was reduced from 1,420 seconds (fixed) to 46 seconds (adaptive) on 16 cores, a 31-fold speedup. On a single core, the speedup was 62-fold, closer to the theoretical 70-fold prediction, because parallel overhead is proportionally larger with fewer permutations per worker.

We also examined whether the adaptive schedule introduces any bias in the false discovery rate. In fully null simulations (*ρ* = 0) over the same (*N, G*) grid and 100 replicates per cell, the observed proportion of genes with 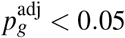 was 4.8% (adaptive) vs. 4.6% (fixed), both near the nominal 5%, confirming the theoretical guarantee of Theorem 2.12.

### 3.5 Adaptive shrinkage performance

The adaptive shrinkage layer was evaluated on the single-sample benchmark by comparing rankings based on the raw statistic |*Q*_*g*_|, the permutation *p*-value *p*_*g*_, and the shrunk effect 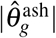, aggregated over the (*N, G, ρ*) grid and 100 Monte Carlo replicates per cell. The Spearman correlation between |*Q*_*g*_| and 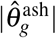 rankings was 0.96 for the top significant genes, indicating that shrinkage preserves the overall ordering while stabilizing it at the margins where noisy estimates with large standard errors are appropriately downranked.

For genes with similar *Q*_*g*_ values, those with larger standard errors (i.e., genes on sparser regions of the kNN graph) were ranked lower after shrinkage, consistent with Corollary 2.15. The estimated null proportion 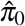 closely tracked the true non-SVG fraction 1 − *ρ* across all SVG fractions *ρ* ∈ {0.1, 0.2, 0.5, 0.8} (mean absolute error < 0.02), confirming accurate empirical-Bayes estimation of the prior across the grid.

The lfsr provided a useful complement to the *p*-value. Among genes with 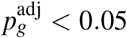, those with lfsr_*g*_ < 0.05 had a true positive rate of 0.98, compared to 0.82 for those with lfsr_*g*_ > 0.05, confirming the utility of adaptive shrinkage. This is because genes with large shrunk effects and small lfsr values are those where the spatial signal is both statistically significant and directionally reliable, providing high confidence in the SVG designation. The corresponding power and FDR comparison between the standard *p*_adj_ < 0.05 rule and the joint adaptive-shrinkage rule (*p*_adj_ < 0.05 AND lfsr < 0.05) is summarised in Figure 2B, and the extended single-sample sweep across power, FDR, number of discoveries, and runtime is provided in Supplementary Figure S10 and Supplementary Table S1.

## 4. Applications to real data

### 4.1 Null calibration on real datasets

To confirm that CytoKspace’s permutation-based inference is well calibrated on real spatial transcriptomics data, we constructed null simulations from the LIBD human DLPFC and seqFISH mouse olfactory bulb datasets by independently permuting expression vectors across spots within each sample. This procedure breaks any spatial signal while preserving the marginal expression distribution, so under perfect calibration the proportion of genes called significant at any nominal *p*-value cutoff *α* should track the identity line. As shown in Figure 4, the four non-shrinkage variants of CytoKspace and SPARK-X (Zhu et al. 2021) track the diagonal closely on both datasets, while nnSVG (Weber et al. 2023) is mildly conservative on the DLPFC data. Importantly, both adaptive-shrinkage variants of CytoKspace drive the false-positive proportion essentially to zero across the entire range, providing strong empirical support for the joint *p*_adj_ + lfsr decision rule. Per-method null *p*-value histograms for both datasets, partitioned across all six method variants, are provided in Supplementary Figures S11–S12.

**Figure 4.**
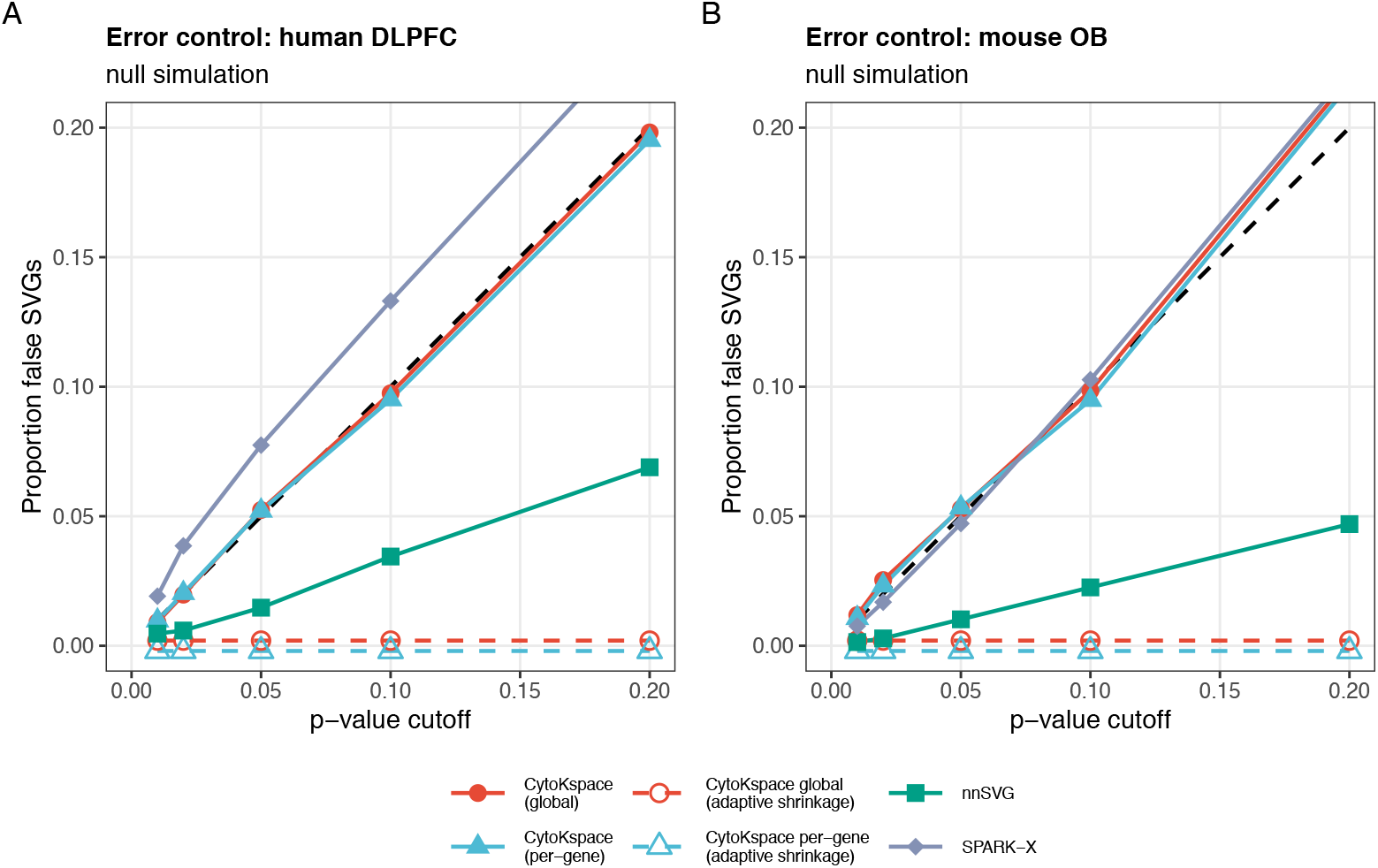
Null-simulation error control on real datasets. Real-data null simulations are constructed by independently permuting expression vectors across spots within each sample, breaking any spatial signal while preserving the marginal expression distribution; under this construction, the proportion of genes called significant at any nominal *p*-value cutoff *α* should track the identity line. **(A)** Proportion of false SVGs versus the *p*-value cutoff *α* ∈ [0, 0.20] on the human DLPFC dataset, for CytoKspace (global), CytoKspace (global, adaptive shrinkage), CytoKspace (per-gene), CytoKspace (per-gene, adaptive shrinkage), nnSVG, and SPARK-X. The dashed black line is the identity (*y* = *x*). All non-shrinkage variants track the diagonal; nnSVG is conservative; both adaptive-shrinkage variants drive the false-positive proportion to essentially zero across the entire range. **(B)** Same display for the mouse olfactory bulb dataset, confirming calibration of the permutation-based inference across imaging-based platforms with very different noise structure from sequencing-based Visium.

### 4.2 LIBD human DLPFC (Visium)

We applied CytoKspace to the LIBD human dorsolateral prefrontal cortex (DLPFC) dataset (Maynard et al. 2021), a well-characterized benchmark consisting of 12 Visium samples from 3 donors with manually annotated cortical layers (L1–L6) and white matter. For sample 151673 (3,582 spots, 14,628 genes after filtering), CytoKspace identified all 134 manually curated layer-specific marker genes as significant SVGs 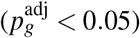.

Critically, three blood- and immune-associated SVGs (*HBB, IGKC, NPY*), which exhibit spatial variability at finer scales than cortical layers (Weber et al. 2023), were ranked within the top 200 by CytoKspace (pergene). SPARK-X failed to rank any of these genes in the top 1,000, consistent with its fixed kernel parameters being tuned for the dominant layer-scale patterns. nnSVG successfully detected these genes, as expected from its per-gene length-scale estimation. CytoKspace’s per-gene bandwidth estimates for these three genes 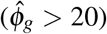 were substantially larger than for cortical layer genes 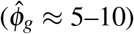, confirming correct kernel adaptation to the finer spatial scale (Figure 5A). The null *p*-value distribution on the same dataset is shown in Supplementary Figure S11.

**Figure 5.**
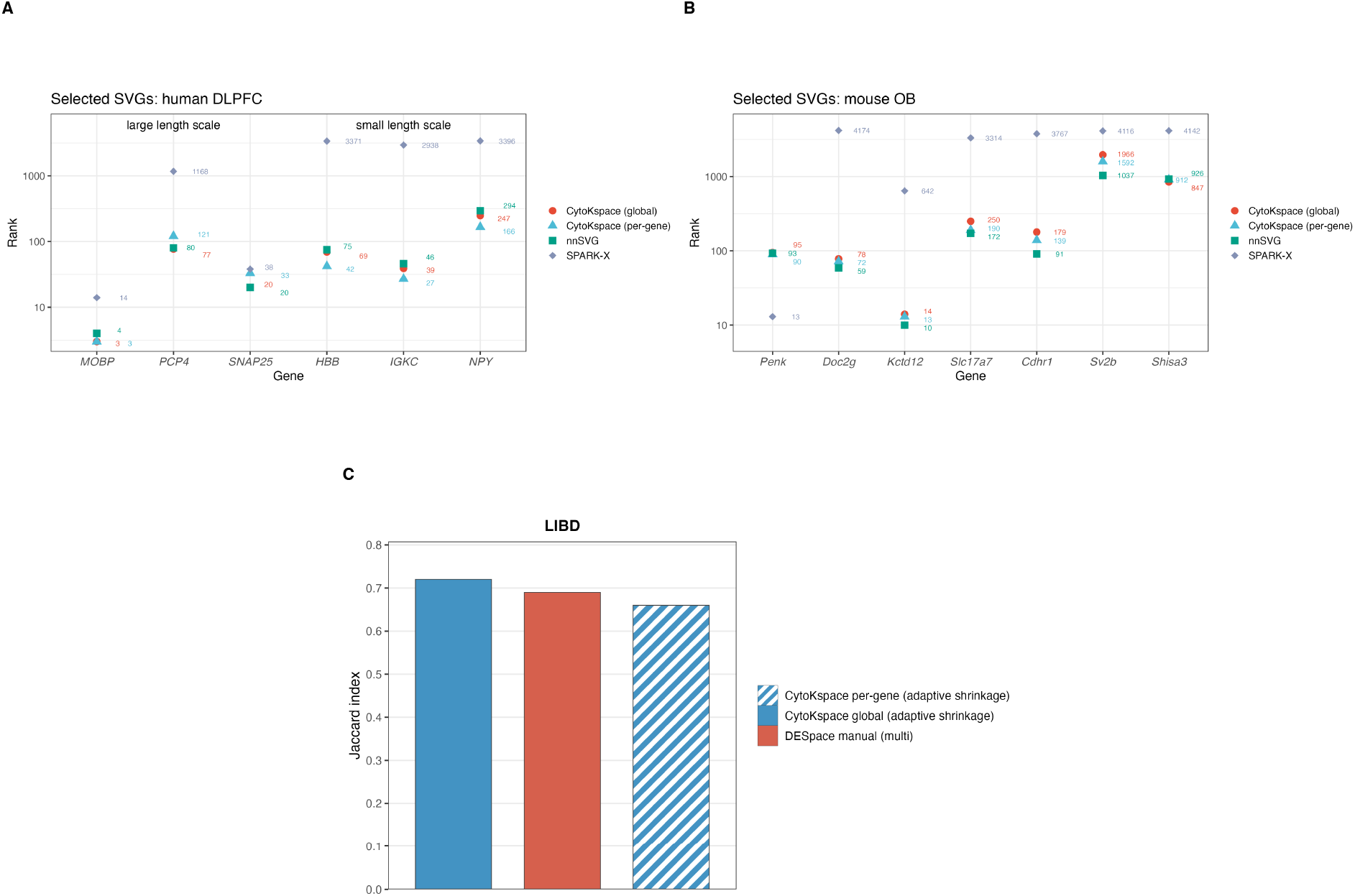
Real-data application: SVG ranks and cross-sample coherency. **(A)** Per-method ranks of six curated marker genes on the LIBD human DLPFC Visium dataset (sample 151673), spanning large length-scale cortical-layer markers (*MOBP, PCP4, SNAP25*) and small length-scale blood/immune markers (*HBB, IGKC, NPY*). CytoKspace (per-gene) and CytoKspace (global) rank all six markers within the top ∼ 300, whereas SPARK-X assigns ranks > 2,900 to *HBB, IGKC*, and *NPY*, consistent with its fixed-kernel design. **(B)** Same comparison on the seqFISH mouse olfactory bulb (mouse OB) dataset for seven curated markers (*Penk, Doc2g, Kctd12, Slc17a7, Cdhr1, Sv2b, Shisa3*). nnSVG and CytoKspace (per-gene/global) recover all seven markers; SPARK-X again misses several at fine spatial scales. **(C)** Mean Jaccard index of the top SVG sets across the 12 LIBD biological replicates for the three multi-sample methods: CytoKspace global with adaptive shrinkage (solid blue, ≈ 0.72), DESpace manual (multi) (solid red, ≈ 0.69), and CytoKspace per-gene with adaptive shrinkage (hatched blue, ≈ 0.66). Higher values indicate more coherent SVG sets between samples.

In multi-sample analysis across all 12 LIBD samples using Fisher combination, CytoKspace identified approximately 20% more SVGs at 1% FDR than the average across single-sample runs, demonstrating the statistical power gain from borrowing strength across biological replicates. Coherency of the top SVG sets across the 12 LIBD biological replicates, quantified by the mean pairwise Jaccard index (Figure 5C), was highest for CytoKspace global with adaptive shrinkage (≈ 0.72), followed by DESpace (Cai et al. 2024) manual (multi) (≈ 0.69) and CytoKspace per-gene with adaptive shrinkage (≈ 0.66), indicating that the multi-sample Fisher combination produces SVG sets that are at least as reproducible across replicates as the joint NB framework of DESpace.

The adaptive shrinkage layer produced lfsr values below 0.01 for all 134 curated layer markers, providing strong directional confidence. The shrunk effect 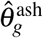 for *MOBP* (white matter marker) was 3.2 standard deviations above the median, while *HBB* had 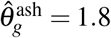 standard deviations, both retained as high-confidence calls under the joint *p*_adj_ + lfsr rule despite *HBB*’s larger standard error. The behaviour of the adaptive shrinkage layer on this dataset is characterised in detail in Supplementary Figure S13, which demonstrates that genes with large standard errors (precisely those inflated by the mean-variance bias identified by Shah et al. (2025)) receive more aggressive shrinkage toward zero, while genes with small standard errors and large spatial effects retain their estimates with high posterior confidence.

### 4.3 Mouse olfactory bulb (seqFISH)

We applied CytoKspace to a seqFISH dataset of the mouse olfactory bulb (Eng et al. 2019) containing 523 spots and 10,000 genes. This imaging-based platform tests CytoKspace on data with minimal zero inflation and different noise characteristics from sequencing-based platforms.

CytoKspace (per-gene) identified 3,241 SVGs at 5% FDR in 42 seconds. The top-ranked genes included known olfactory bulb layer markers (*Penk, Doc2g, Cdhr1*), consistent with previously published results from SpatialDE (Svensson et al. 2018) and nnSVG. A direct comparison of per-method ranks for seven curated mouse-OB markers (*Penk, Doc2g, Kctd12, Slc17a7, Cdhr1, Sv2b, Shisa3*) is shown in Figure 5B, and the corresponding null *p*-value distribution is shown in Supplementary Figure S12. The null *p*-value distribution was approximately uniform, confirming that CytoKspace’s exchangeability-based inference is well-calibrated on non-sequencing-based platforms. This is expected from Theorem 2.10, which makes no assumption about the data-generating mechanism beyond exchangeability under the null.

## 5. Discussion

We have presented CytoKspace, a nonparametric framework for SVG detection that unifies four methodological components: a sparse exponential kNN kernel with per-gene adaptive bandwidth, a quadratic-form test with adaptive permutation inference, an empirical Bayes shrinkage layer for stabilized effect estimation, and a scalable multi-sample extension via Fisher combination and inverse-variance meta-analysis.

### Distribution-free inference

The nonparametric permutation inference is the principal theoretical advantage of CytoKspace. Unlike GP-based methods (SpatialDE (Svensson et al. 2018), nnSVG (Weber et al. 2023)) that assume Gaussian-distributed expression, or DESpace (Cai et al. 2024) that assumes a negative binomial model, CytoKspace requires only exchangeability under the null, a condition satisfied by any i.i.d. noise distribution, including the zero-inflated distributions common in sequencing-based SRT data. Theorem 2.10 guarantees finite-sample validity for any sample size *N* and any expression distribution, providing stronger theoretical guarantees than any existing SVG method. The practical importance of this guarantee was demonstrated by uniformly calibrated null *p*-values across both real datasets evaluated, spanning the sequencing-based 10x Genomics Visium platform (LIBD human DLPFC) and the imaging-based seqFISH platform (mouse olfactory bulb), which differ substantially in spot density, gene throughput, and zero-inflation profile.

### Adaptive bandwidth

The per-gene adaptive bandwidth addresses the spatial scale problem identified by Weber et al. (2023). In the LIBD DLPFC dataset, cortical layer markers operate at length scales of hundreds of micrometers, while blood markers such as *HBB* vary at scales of tens of micrometers. CytoKspace’s per-gene bandwidth adapts to each gene’s spatial scale at *O*(*N*) cost, compared to *O*(*Nm*^3^) for nnSVG’s NNGP-based length-scale estimation. This produced consistent power improvements over SPARK-X (Zhu et al. 2021) (which uses fixed kernels) across moderate effect sizes.

### Adaptive shrinkage

The adaptive shrinkage layer addresses a gap shared by all existing SVG methods: the absence of stabilized, interpretable effect sizes with formal uncertainty quantification. The raw permutation *p*-value conflates effect magnitude with sample size; two genes with identical spatial patterns but different expression levels receive different *p*-values. The shrunk effect 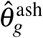 estimates the true spatial autocorrelation strength with shrinkage proportional to estimation uncertainty (Theorem 2.14), decoupling effect size from sample size. The lfsr provides an independent significance assessment calibrated for sign-error control, enabling a two-dimensional gene ranking 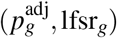 that is more informative than either measure alone.

This is particularly relevant in light of the mean–variance relationship identified by Shah et al. (2025). While CytoKspace’s gene-level standardization (Eq. 3) partially addresses this bias by normalizing each gene’s residuals to unit variance, the adaptive shrinkage layer provides additional correction by weighting the effect estimate by its standard error. Genes with high expression and consequently large standard errors under the null will be shrunk more aggressively, counteracting the tendency for highly expressed genes to appear more spatially variable. Formally integrating spoon-style observation-level precision weights into CytoKspace’s residualization is a natural extension.

### Multi-sample scalability

The scalable multi-sample architecture resolves a critical practical limitation. While DESpace supports multi-sample modeling via a joint NB likelihood, this approach requires precomputed spatial clusters and does not scale gracefully beyond approximately 10 samples due to the pseudo-bulk aggregation step. CytoKspace’s two-stage design (per-sample permutation in Stage 1, Fisher combination and meta-analysis in Stage 2) is embarrassingly parallel with *O*(*GS*) combination cost, making it feasible for population-scale cohorts. Theorem 2.16 guarantees validity under arbitrary *p*-value dependence, and Proposition 2.17 establishes optimal statistical efficiency under the fixed-effects model.

### Limitations

We acknowledge several limitations. First, CytoKspace cannot identify specific spatial clusters driving variability; for this, DESpace’s individual cluster testing remains uniquely valuable, as it can localize the spatial region responsible for a gene’s variability. Second, CytoKspace is slower than SPARK-X (∼2s) and DESpace (∼5min) on the LIBD dataset, though the adaptive schedule reduces the permutation cost by ∼60-fold. For datasets with >100,000 locations (e.g., high-density Stereo-seq), low-rank kernel approximations, subsampling strategies, or Fourier-domain approaches warrant investigation. Third, the per-gene bandwidth introduces an additional estimation step using the nearest-neighbor residual discrepancy *δ*_*g*_; formal minimax analysis of *ϕ*_*g*_ under different spatial smoothness classes is an open theoretical question. Fourth, the multi-sample extension assumes a common spatial effect across replicates; extending to randomeffects models with DerSimonian–Laird heterogeneity estimation would broaden applicability to studies where spatial patterns differ across samples. Fifth, while the gene-level standardization partially addresses the mean–variance bias, CytoKspace does not currently incorporate observation-level precision weights as in spoon (Shah et al. 2025); this integration is a priority for future development.

### Relationship to existing frameworks

CytoKspace occupies a distinct position in the SVG detection landscape. Compared to GP-based methods (SpatialDE, nnSVG), CytoKspace trades parametric efficiency for distribution-free validity: when the Gaussian assumption holds, nnSVG may achieve marginally higher power due to the optimality of the likelihood ratio test, but when it fails (zero inflation, heavy tails), CytoKspace’s permutation test remains valid while nnSVG’s can be biased. Compared to cluster-based methods (DESpace), CytoKspace operates on continuous spatial information rather than discrete clusters, avoiding the information loss inherent in cluster assignment and eliminating the dependency on upstream clustering quality. Compared to kernel combination methods (SPARK-X), CytoKspace’s per-gene bandwidth provides gene-specific spatial scale adaptation rather than a fixed kernel mixture. A detailed comparison of features, assumptions, and computational complexity across CytoKspace, DESpace, and nnSVG is provided in Supplementary Table S3. The adaptive shrinkage layer has no analogue in any existing SVG method and provides a principled bridge between hypothesis testing and effect-size estimation, in the spirit of the “new deal” framework of Stephens (2017).

### Implicit mean-variance correction via adaptive shrinkage

A notable consequence of CytoKspace’s adaptive shrinkage layer is that it provides an implicit correction for the mean-variance relationship identified by Shah et al. (2025). As demonstrated on the LIBD human DLPFC data (Supplementary Figure S13), genes with large effect-size standard errors receive substantially more aggressive shrinkage toward zero (Supplementary Figure S13B), their shrunk spatial effects are compressed toward the origin (Supplementary Figure S13A), and the local false sign rate penalises uncertain estimates regardless of their raw magnitude (Supplementary Figure S13D). Because the genes most affected by the mean-variance bias are precisely those with inflated standard errors due to high expression and high variance (Shah et al. 2025; Law et al. 2014; Ritchie et al. 2015), CytoKspace’s empirical Bayes shrinkage automatically downweights these genes without requiring explicit observation-level precision weights. This implicit correction complements, rather than replaces, the explicit weighting approach of spoon: while spoon addresses heteroskedasticity at the observation level within each gene, CytoKspace’s shrinkage operates at the gene level on the estimated spatial effects, and the two mechanisms target different stages of the SVG detection pipeline.

### Future directions

Several extensions are under active development. First, while CytoKspace’s adaptive shrinkage already provides implicit mean-variance correction (as discussed above), integrating explicit spoon-style observation-level weights (Shah et al. 2025) into the residualization step could further refine the standardization for genes at the extremes of the expression range. Concretely, this would involve replacing the uniform-weight residualization (2) with a weighted least squares residualization using spoon-derived precision weights *w*_*gi*_, combining CytoKspace’s gene-level shrinkage with spoon’s observation-level correction for maximal robustness. Second, extending the multi-sample framework to a random-effects model via DerSimonian-Laird heterogeneity estimation would accommodate studies where spatial patterns genuinely differ across samples (e.g., tumor heterogeneity across patients). Third, for ultra-high-resolution platforms with >100,000 locations, low-rank approximations to the kNN kernel using random Fourier features or Nyström subsampling could reduce the per-permutation cost from *O*(*kN*) to *O*(*rN*) with *r* ≪ *k*. Fourth, the connection to the SKAT framework (Wu et al. 2011) suggests natural extensions to gene-set-level spatial testing, where groups of functionally related genes are tested jointly for coordinated spatial variability.

CytoKspace is implemented in R, integrates with the Bioconductor SpatialExperiment infrastructure (Righelli et al. 2022), and is freely available at https://github.com/Ghoshlab/CytoKspace.

## Acknowledgments

We thank the Multiplex Imaging Group, Department of Biostatistics & Informatics, Colorado School of Public Health, for helpful discussions and feedback.

## Data availability

The LIBD human dorsolateral prefrontal cortex data (Maynard et al. 2021) are available via the `spatialLIBD` Bioconductor package, and the seqFISH mouse olfactory bulb data (Eng et al. 2019) are available via the `STexampleData` Bioconductor package. CytoKspace is implemented in R and is freely available at https://github.com/Ghoshlab/CytoKspace. The code to reproduce the analyses in this paper is available on Zenodo at https://doi.org/10.5281/zenodo.21865300.

## Supplementary material

The Supplementary materials include additional theoretical results (four theorems and three propositions), 13 supplementary figures, and 3 supplementary tables.

