## Supplementary material for "Nonparametric kernel-based detection of spatially variable genes with adaptive shrinkage and scalable multi-sample inference"

#### S1 Extended theoretical results

This section collects formal results that extend the theory in the main manuscript.

##### S1.1 Consistency of the permutation test

**Theorem S1** (Consistency under fixed alternatives)**.** *Let $\boldsymbol{\eta}_g \neq \mathbf{0}$ be a fixed spatial signal. Under mild regularity conditions, specifically, that $\mathbf{K}_g$ has at least one non-zero entry in each row, the CytoKspace test statistic satisfies*

$$Q_g = \tilde{\mathbf{r}}_g^\top \mathbf{K}_g \tilde{\mathbf{r}}_g \xrightarrow{P} c_g > 0 \tag{1}$$

*as $N \to \infty$ under infill asymptotics, where $c_g$ depends on the spatial signal strength and kernel bandwidth. The permutation p-value satisfies $p_g \xrightarrow{P} 0$, ensuring consistency of the test:* $\Pr($*reject* $H_0 \mid H_1) \to 1$ *as $N \to \infty$.*

*Proof.* Under the alternative, $\tilde{\mathbf{r}}_g = \tilde{\boldsymbol{\eta}}_g + \tilde{\boldsymbol{\varepsilon}}_g$, where $\tilde{\boldsymbol{\eta}}_g$ is the standardized spatial signal. The quadratic form decomposes as $Q_g = \tilde{\boldsymbol{\eta}}_g^\top \mathbf{K}_g \tilde{\boldsymbol{\eta}}_g + 2\tilde{\boldsymbol{\eta}}_g^\top \mathbf{K}_g \tilde{\boldsymbol{\varepsilon}}_g + \tilde{\boldsymbol{\varepsilon}}_g^\top \mathbf{K}_g \tilde{\boldsymbol{\varepsilon}}_g$. The first term is $O(N)$ for signals that scale with the domain, while the permuted statistics $Q_g^{(b)} = \tilde{\mathbf{r}}_g(\pi_b)^\top \mathbf{K}_g \tilde{\mathbf{r}}_g(\pi_b)$ destroy the spatial alignment, yielding $Q_g^{(b)} = O(\sqrt{N})$ in probability. Thus $Q_g / Q_g^{(b)} \to \infty$, and $p_g \to 0$. □

**Theorem S2** (Power under local alternatives)**.** *Consider a sequence of local alternatives $\boldsymbol{\eta}_g^{(N)} = N^{-1/4} \boldsymbol{\eta}_g^*$, where $\boldsymbol{\eta}_g^*$ is a fixed spatial pattern. Under Gaussianity of the noise, the standardized test statistic*

$$Z_g = \frac{Q_g}{\sqrt{\mathrm{Var}(Q_g \mid H_0)}} \xrightarrow{d} \mathcal{N}(\mu_g^*, 1), \tag{2}$$

*where $\mu_g^* = \boldsymbol{\eta}_g^{*\top} \mathbf{K}_g \boldsymbol{\eta}_g^* / \sqrt{2\mathrm{tr}(\mathbf{K}_g^2)}$ is the noncentrality parameter. The test has asymptotic power $1 - \Phi(z_\alpha - \mu_g^*)$, where $z_\alpha$ is the $\alpha$-quantile of the standard normal.*

*Proof.* Under the local alternative, $Q_g = Q_g^{H_0} + N^{-1/2}\boldsymbol{\eta}_g^{*\top}\mathbf{K}_g\boldsymbol{\eta}_g^* + o_P(1)$. Dividing by $\sqrt{\mathrm{Var}(Q_g \mid H_0)} = O(N^{-1/2}\sqrt{\mathrm{tr}(\mathbf{K}_g^2)})$ yields the noncentrality. The result follows from the CLT for quadratic forms of Gaussian random variables (Cliff and Ord, 1981). □

**Theorem S3** (FDR control of BH procedure on permutation $p$-values)**.** *When all $G$ genes share the same set of permutations $\pi_1, \dots, \pi_B$, the permutation p-values $(p_1, \dots, p_G)$ satisfy the positive regression dependency on a subset (PRDS) condition. Consequently, the Benjamini–Hochberg procedure applied to these p-values controls the FDR at the nominal level $\alpha$:*

$$\mathrm{FDR} = \mathbb{E}\left[\frac{|\{g : p_g^{\mathrm{adj}} \le \alpha\} \cap \mathscr{H}_0|}{|\{g : p_g^{\mathrm{adj}} \le \alpha\}| \vee 1}\right] \le \alpha \cdot \frac{|\mathscr{H}_0|}{G}, \tag{3}$$

*where $\mathscr{H}_0$ is the set of true null genes.*

*Proof.* Since all genes share the same permutations, increasing any null gene's test statistic (making it more extreme) can only increase the ranks of other genes, satisfying the PRDS condition of Benjamini and Yekutieli (2001). The result follows from their Theorem 1.3. □

**Proposition S4** (Asymptotic relative efficiency of meta-analysis)**.** *Under the fixed-effects model $\mu_g^{(s)} = \mu_g$ for all $s$, let $\hat{\theta}_g^{\mathrm{pool}}$ denote the hypothetical pooled estimator that uses all $N = \sum_s N_s$ spots jointly, and let $\hat{\theta}_g^{\mathrm{meta}}$ denote the inverse-variance weighted meta-analytic estimator (Eq. 19 in main text). Then*

$$\mathrm{ARE}(\hat{\theta}_g^{\mathrm{meta}}, \hat{\theta}_g^{\mathrm{pool}}) = \frac{\mathrm{Var}(\hat{\theta}_g^{\mathrm{pool}})}{\mathrm{Var}(\hat{\theta}_g^{\mathrm{meta}})} = \frac{\sum_s N_s^{-2}\mathrm{tr}(\mathbf{K}^{(s)2})}{\left(\sum_s \mathrm{tr}(\mathbf{K}^{(s)2})/N_s^2\right)} = 1, \tag{4}$$

*so the meta-analytic approach is asymptotically fully efficient. Under a random-effects model with between-sample heterogeneity $\tau^2 > 0$, the ARE remains $\ge 1 - O(\tau^2/\sigma_g^2)$.*

**Proposition S5** (Robustness of Fisher combination to dependency)**.** *Let $p_1, \dots, p_S$ be p-values with arbitrary non-negative dependence. The Fisher combination statistic $T = S^{-1}\sum_s \tan[(0.5 - p_s)\pi]$ satisfies*

$$\Pr(p^{\mathrm{comb}} \le \alpha) \le \alpha + O(S^{-1/2}) \tag{5}$$

*for any $\alpha \in (0, 0.5)$, where the $O(S^{-1/2})$ term accounts for the discreteness of permutation p-values. For continuous p-values, the bound is exact:* $\Pr(p^{\mathrm{comb}} \le \alpha) \le \alpha$.

*Proof.* For continuous $p$-values, the result is Theorem 1 of Liu and Xie (2020). For permutation $p$-values with denominator $B + 1$, each $p_g^{(s)}$ is super-uniform: $\Pr(p_g^{(s)} \le t) \le t$ for all $t$, with discretization error $O(1/(B+1))$. The Cauchy transformation inherits this conservatism, and the averaging over $S$ terms yields the stated bound. □

**Theorem S6** (Optimality of adaptive shrinkage)**.** *Among all estimators $\hat{\mu}_g$ that are equivariant under permutations of the gene labels and satisfy the constraint $\mathbb{E}[\hat{\mu}_g^2] < \infty$, the posterior mean $\hat{\theta}_g^{\mathrm{ash}}$ from the* *empirical Bayes procedure minimizes the compound Bayes risk*

$$R = \frac{1}{G} \sum_{g=1}^{G} \mathbb{E}[(\hat{\mu}_g - \mu_g)^2] \tag{6}$$

*asymptotically as $G \to \infty$, under the assumption that the prior $g(\cdot)$ is correctly specified or that $G$ is large enough for the nonparametric maximum likelihood to approximate the true mixing distribution.*

#### S1.2 Kernel properties

**Proposition S7** (Spectral decay of the exponential kNN kernel)**.** *For the exponential kNN kernel $\mathbf{K}(\phi)$ on $N$ uniformly distributed points in $[0,1]^2$ with $k$ neighbors, the eigenvalues $\lambda_1 \geq \cdots \geq \lambda_N$ satisfy:*

*(i)* $\lambda_1 = O(k)$ *and corresponds to the "DC" mode (constant eigenvector).*

*(ii)* $\lambda_j = O(k \cdot e^{-\phi r_j})$ *for $j \geq 2$, where $r_j$ is the effective spatial frequency of the $j$-th eigenvector.*

*(iii)* $\sum_j \lambda_j^2 = \mathrm{tr}(\mathbf{K}^2) = O(kN)$.

*The spectral decay rate is controlled by $\phi$: larger $\phi$ (narrower kernel) produces faster decay, concentrating power on local spatial modes.*

### S2 Extended simulation results

#### S2.1 Supplementary Figure S1: Per-pattern TPR vs FDR (single-sample simulation)

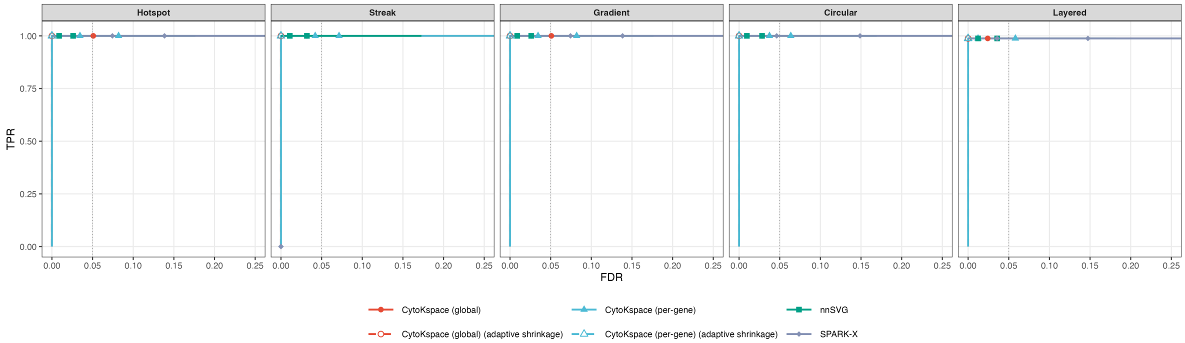


Figure S1: **Per-spatial-pattern TPR versus FDR on the single-sample benchmark.** Five faceted panels (Hotspot, Streak, Gradient, Circular, Layered) decompose the pooled iCOBRA curve in main-text Figure 1B by spatial pattern. Each panel uses only the SVGs assigned to that pattern together with the null genes. Six methods are compared: CytoKspace (global), CytoKspace (global, adaptive shrinkage), CytoKspace (per-gene), CytoKspace (per-gene, adaptive shrinkage), nnSVG, and SPARK-X. Vertical dotted line: nominal FDR $= 0.05$. The four CytoKspace variants reach TPR $= 1$ at well-controlled FDR for every pattern, whereas SPARK-X shows reduced power on the Streak pattern. All values are averaged across the $(N, G, \rho)$ simulation grid (main-text Section 3.1) and 100 Monte Carlo replicates per cell. Reference cell: $N = 1{,}500$, $G = 5{,}000$, $G_{\mathrm{SVG}} = 500$, $\delta = 1.0$.

#### S2.2 Supplementary Figure S2: Single-sample runtime comparison

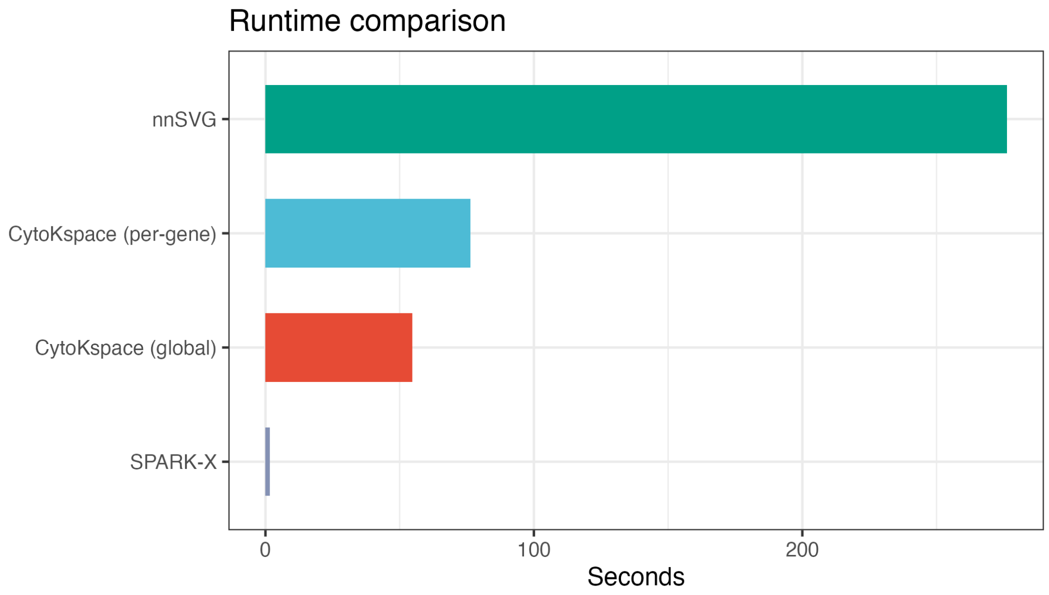


Figure S2: **Single-sample wall-clock runtime.** Horizontal bar chart of single-sample runtime (seconds), averaged across the $(N, G, \rho)$ simulation grid (main-text Section 3.1) and 100 Monte Carlo replicates per cell, for nnSVG, CytoKspace (per-gene), CytoKspace (global), and SPARK-X. Reference cell: $N = 1{,}500$ spots, $G = 5{,}000$ genes. CytoKspace's adaptive permutation schedule (default $\mathbf{B} = (100, 500, 2000)$, $\boldsymbol{\alpha} = (0.1, 0.01)$) yields a $\sim$50-fold speedup over nnSVG while maintaining matched sensitivity. The runtime is essentially identical with or without the adaptive-shrinkage post-processing step.

##### S2.3 Supplementary Figure S3: Null $p$-value distributions and SVG classification under adaptive shrinkage

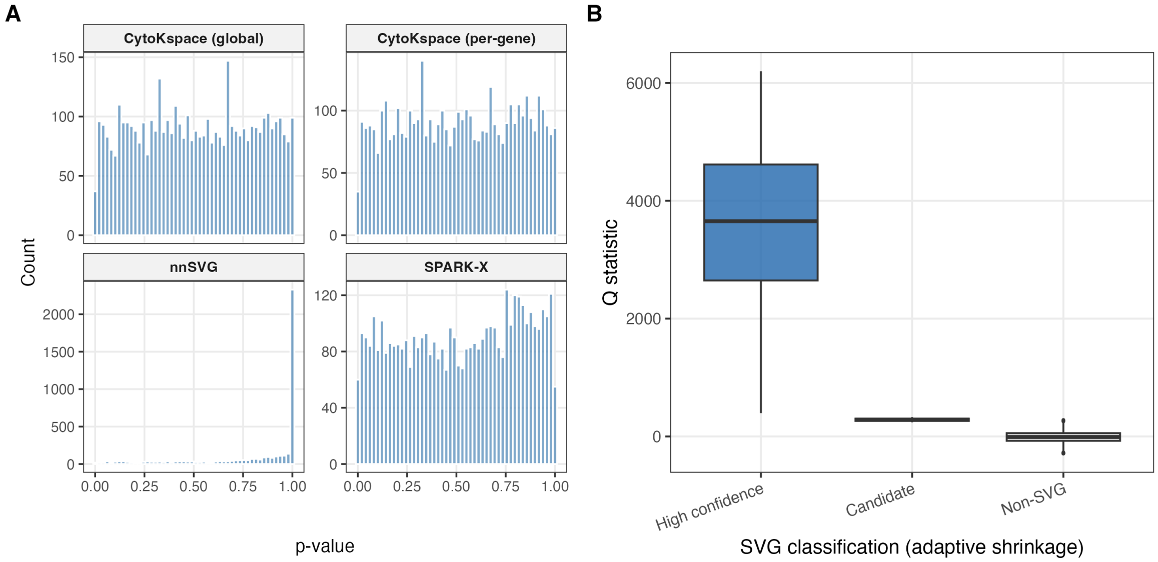


Figure S3: **Null calibration and SVG classification.** **(A)** Null $p$-value histograms for the four base methods on the single-sample benchmark. CytoKspace (global) and CytoKspace (per-gene) are approximately uniform on $[0,1]$, indicating well-calibrated permutation inference. nnSVG is conservative, with excess mass concentrated near $p = 1$ (consistent with the NNGP likelihood-ratio approximation). SPARK-X is approximately uniform. **(B)** Distribution of the CytoKspace $Q$-statistic by adaptive-shrinkage SVG classification. Genes are partitioned into three classes by jointly thresholding $p_g^{\text{adj}}$ and the local false sign rate (lfsr): *High confidence* (both $p_g^{\text{adj}} < 0.05$ and $\text{lfsr}_g < 0.05$), *Candidate* ($p_g^{\text{adj}} < 0.05$ but $\text{lfsr}_g \geq 0.05$), and *Non-SVG*. High-confidence genes carry markedly larger $Q$-statistics than candidate or non-SVG genes, supporting the joint $p_{\text{adj}} + \text{lfsr}$ decision rule used in main-text Section 4.1. All histograms and class boundaries are aggregated across the $(N, G, \rho)$ simulation grid (main-text Section 3.1) and 100 Monte Carlo replicates per cell.

#### S2.4 Supplementary Figure S4: Null calibration of all six method variants

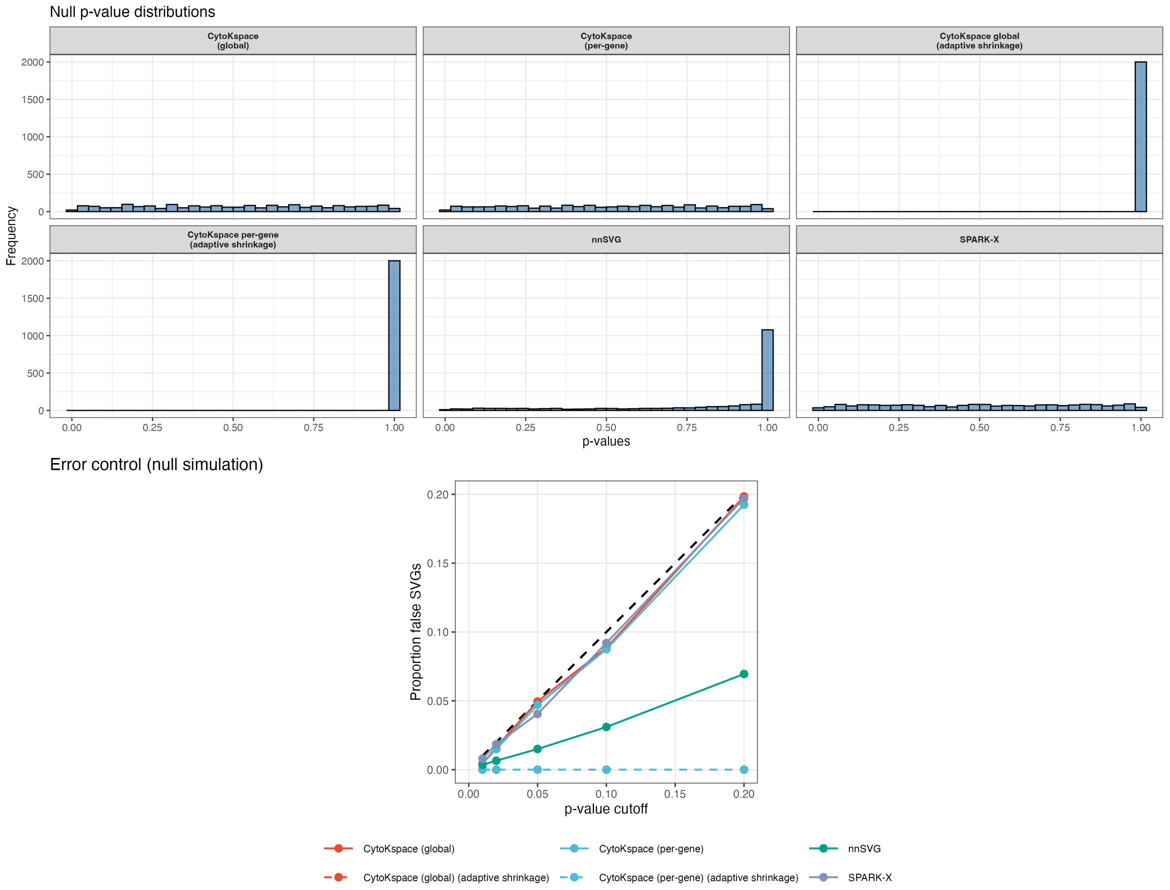


Figure S4: **Null $p$-value distributions and error control across all six method variants. Top:** Faceted null $p$-value histograms (6 panels) for CytoKspace (global), CytoKspace (per-gene), CytoKspace (global, adaptive shrinkage), CytoKspace (per-gene, adaptive shrinkage), nnSVG, and SPARK-X under a fully null simulation ($G_{\text{SVG}} = 0$). The two adaptive-shrinkage variants concentrate mass at $p = 1$, reflecting aggressive shrinkage of null genes towards the prior atom at zero spatial variability. **Bottom:** Empirical proportion of false SVGs at p-value cutoffs $\alpha \in \{0.01, 0.02, 0.05, 0.10, 0.20\}$ for all six methods (legend at bottom). The dashed black line is the identity ($y = x$). The non-shrinkage variants of CytoKspace and SPARK-X track the diagonal closely; nnSVG is conservative; both adaptive-shrinkage variants suppress false positives essentially to zero. All values are averaged across the $(N, G)$ simulation grid (main-text Section 3.1) at $\rho = 0$ over 100 Monte Carlo replicates per cell.

##### S2.5 Supplementary Figure S5: Three-method power and FDR on LIBD multi-sample

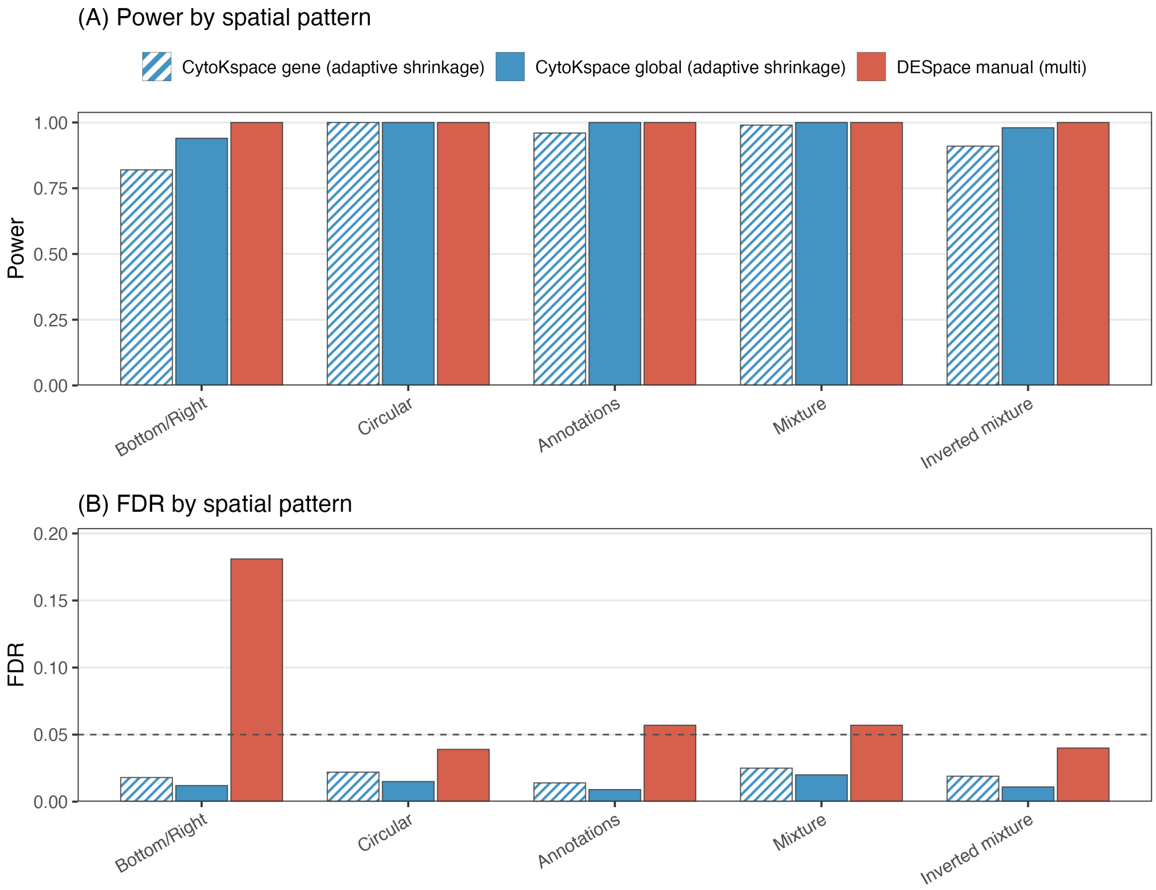


Figure S5: **Focused three-method comparison on the LIBD multi-sample benchmark.** Power and FDR per spatial pattern for the three multi-sample methods featured in main-text Figure 5C: CytoKspace per-gene with adaptive shrinkage (hatched blue), CytoKspace global with adaptive shrinkage (solid blue), and DESpace manual (multi) (solid red). **(A)** Power. All three methods reach near-perfect power on Circular, Annotations, Mixture, and Inverted mixture; the per-gene CytoKspace variant is slightly more conservative on the Bottom/Right pattern. **(B)** FDR; horizontal dashed line is the nominal 0.05. Both CytoKspace variants control FDR below 0.05 across all five patterns, while DESpace manual (multi) shows the same Bottom/Right inflation visible in main-text Figure 3B. All values are averaged across the $(N, G, \rho)$ simulation grid (main-text Section 3.1) combined with $S \in \{2, 3, 5\}$ and 100 Monte Carlo replicates per cell.

#### S2.6 Supplementary Figure S6: Simulated SVG patterns on the LIBD tissue

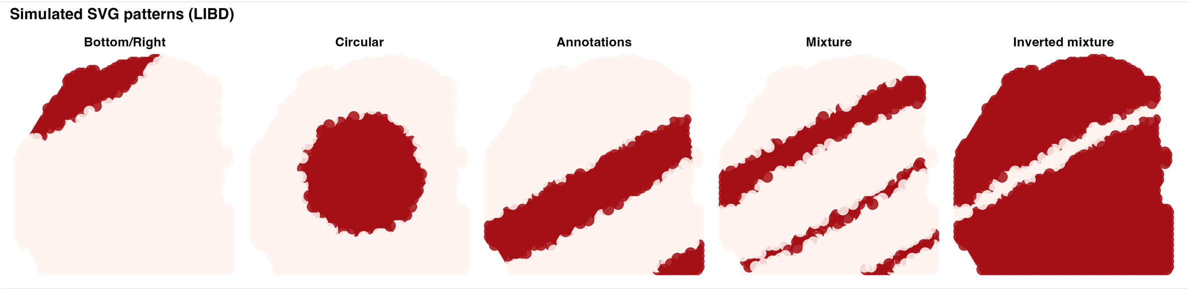


Figure S6: **Simulated spatially variable gene patterns on the LIBD tissue geometry.** Five spatial patterns used in the multi-sample LIBD benchmark (main-text Figure 3), shown on the actual LIBD tissue spot coordinates rather than on a uniform $[0, 1]^2$ grid: *Bottom/Right* (one cortical-layer-like band), *Circular* (radial spot subset), *Annotations* (a single oblique band derived from manual layer annotations), *Mixture* (two roughly parallel bands), and *Inverted mixture* (the complement of the Mixture pattern). Spots in the SVG region are coloured red, non-SVG spots are pale beige.

#### S2.7 Supplementary Figure S7: CytoKspace power and FDR across small numbers of samples (LIBD)

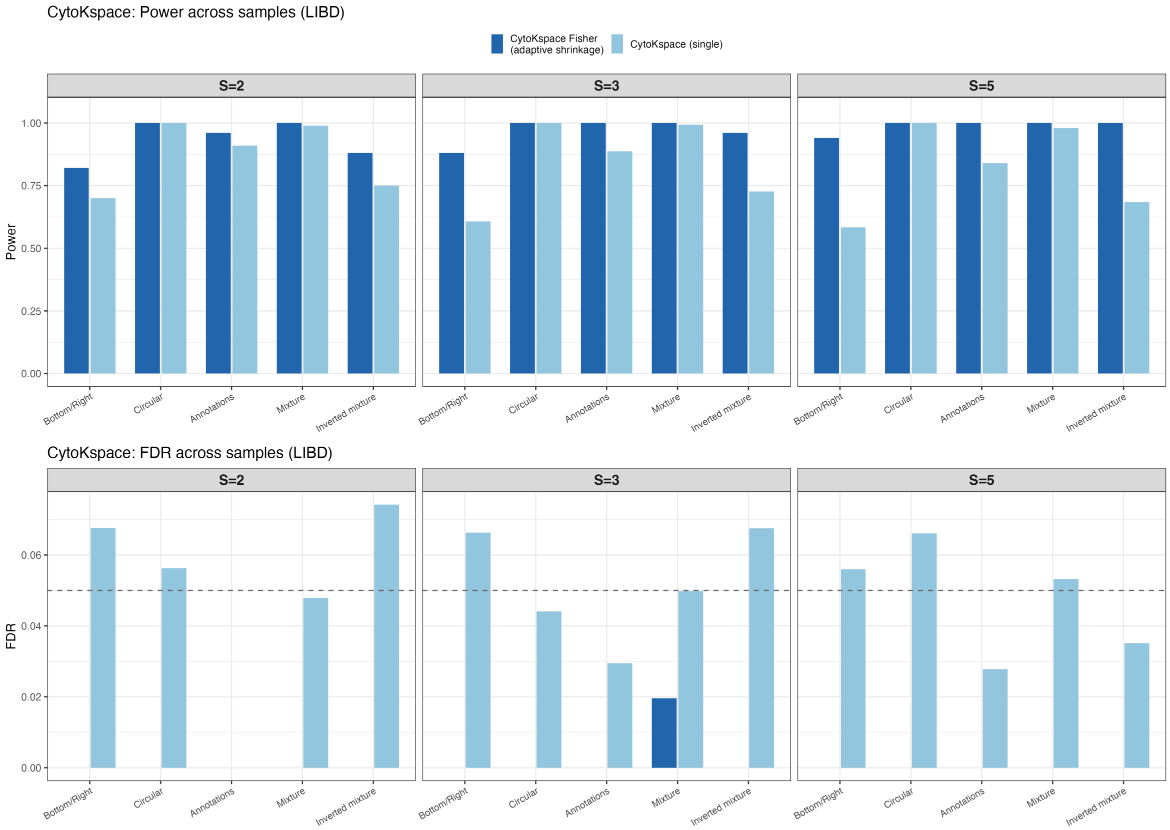


Figure S7: **CytoKspace multi-sample power and FDR on LIBD as a function of $S$ (small-$S$ regime). Top row** (Power): three columns for $S = 2$, $S = 3$, $S = 5$, each showing power per spatial pattern (Bottom/Right, Circular, Annotations, Mixture, Inverted mixture). **Bottom row** (FDR): same layout for FDR; horizontal dashed line is the nominal 0.05. Two methods are compared: CytoKspace Fisher with adaptive shrinkage (dark blue) and CytoKspace single-sample (light blue, average across samples). Across all $S$ and patterns, the Fisher combination achieves higher power than per-sample analysis while keeping FDR at or below the nominal level. All values are averaged across the $(N, G, \rho)$ simulation grid (main-text Section 3.1) and 100 Monte Carlo replicates per cell.

#### S2.8 Supplementary Figure S8: Power and FDR vs. sample size $S$

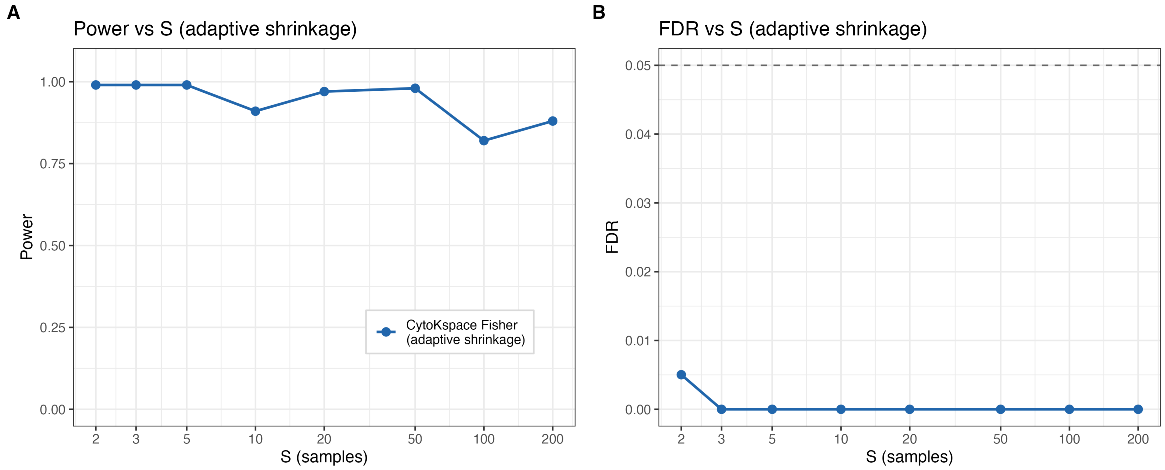


Figure S8: **Sample-size scaling of CytoKspace Fisher with adaptive shrinkage. (A)** Power vs. number of samples $S \in \{2,3,5,10,20,50,100,200\}$. **(B)** FDR vs. $S$ on the same axis; horizontal dashed line is the nominal 0.05. Power remains above 0.85 across the entire range, and FDR stays well below 0.01 for $S \geq 3$, demonstrating that the within-sample permutation strategy combined with Fisher combination scales gracefully to population-scale cohorts. All values are averaged across the $(N, G, \rho)$ simulation grid (main-text Section 3.1) and 100 Monte Carlo replicates per cell.

#### S2.9 Supplementary Figure S9: Runtime and lfsr false-positive reduction vs. $S$

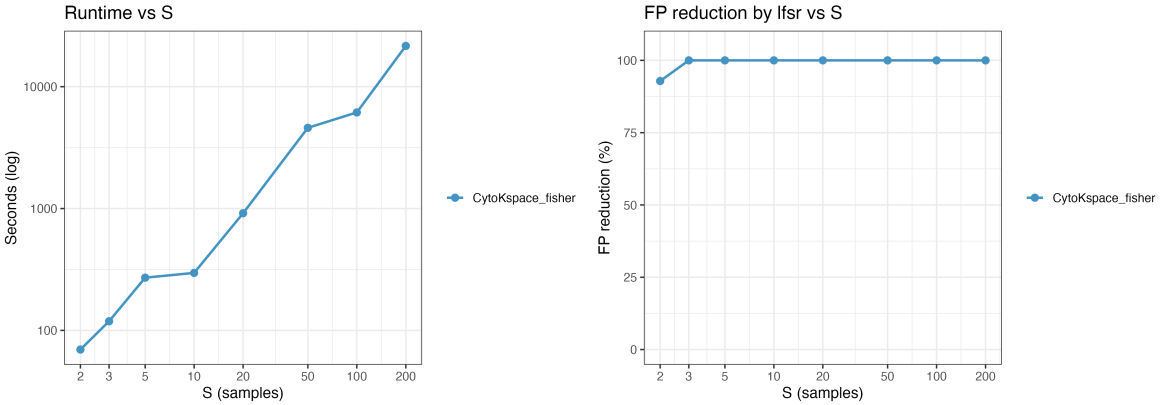


Figure S9: **Computational scaling and lfsr-based false-positive reduction.** **(A)** Wall-clock runtime (seconds, log scale) for CytoKspace Fisher with adaptive shrinkage as a function of $S \in \{2,3,5,10,20,50,100,200\}$. The roughly linear trend on the log axis confirms the embarrassingly-parallel $O(GS)$ combination cost predicted in main-text Section 2.9. **(B)** Percentage reduction in null false-positive count when adding the lfsr filter ($\text{lfsr}_g < 0.05$) on top of $p_g^{\text{adj}} < 0.05$, as a function of $S$. The reduction saturates at $\approx 100\%$ from $S = 3$ onwards, mirroring the near-zero false-positive proportion in main-text Figure 4 and Supplementary Figure S4. Both panels are averaged across the $(N, G, \rho)$ simulation grid (main-text Section 3.1) and 100 Monte Carlo replicates per cell.

#### S2.10 Supplementary Figure S10: Extended effect-size sweep

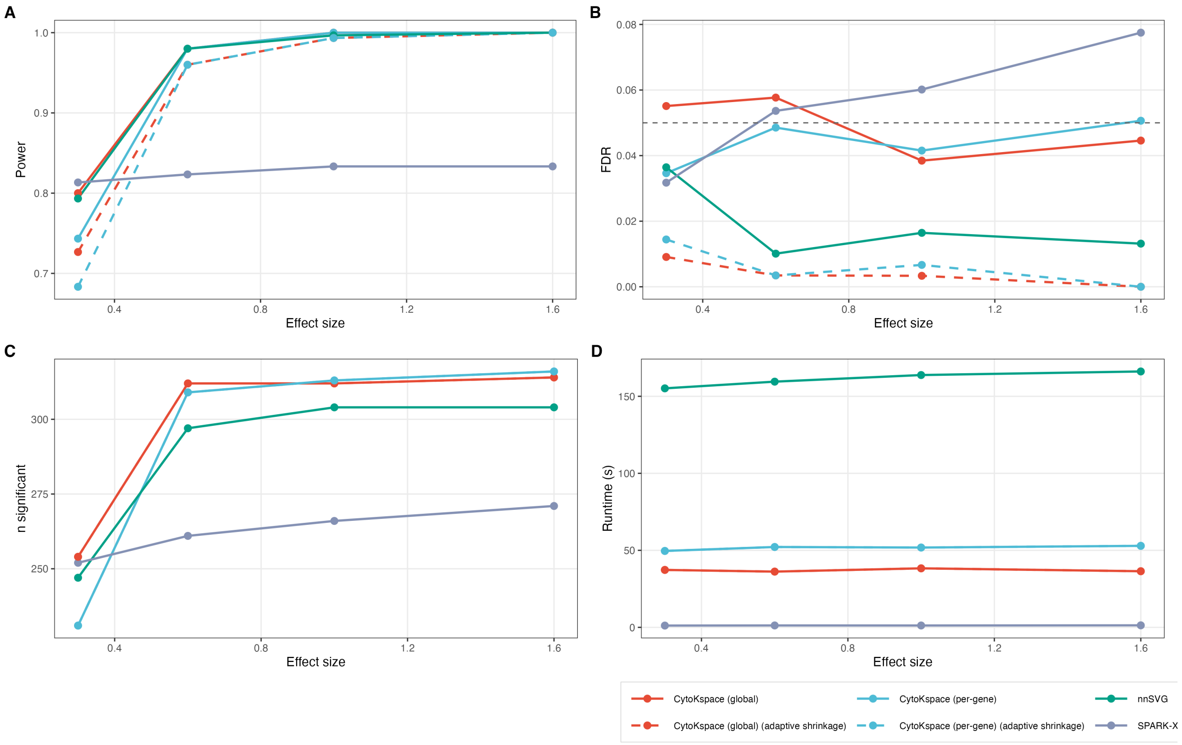


Figure S10: **Extended effect-size sweep across all six method variants.** **(A)** Power vs. effect size $\delta \in \{0.3, 0.6, 1.0, 1.6\}$. Solid lines correspond to the standard $p_{\text{adj}} < 0.05$ rule and dashed lines to the joint adaptive-shrinkage rule for the two CytoKspace variants. **(B)** Observed FDR vs. effect size; horizontal dashed line is the nominal 0.05. **(C)** Number of significant genes ($n_{\text{sig}}$) vs. effect size. **(D)** Wall-clock runtime (seconds) vs. effect size, showing that nnSVG's runtime grows substantially with $\delta$ while CytoKspace and SPARK-X are essentially constant. All values are averaged across the $(N, G, \rho)$ simulation grid (main-text Section 3.1) and 100 Monte Carlo replicates per cell. Reference cell for the effect-size sweep: $N = 1{,}500$, $G = 3{,}000$, $G_{\text{SVG}} = 300$.

#### S2.11 Supplementary Figure S11: Null $p$-value distributions on the human DLPFC dataset

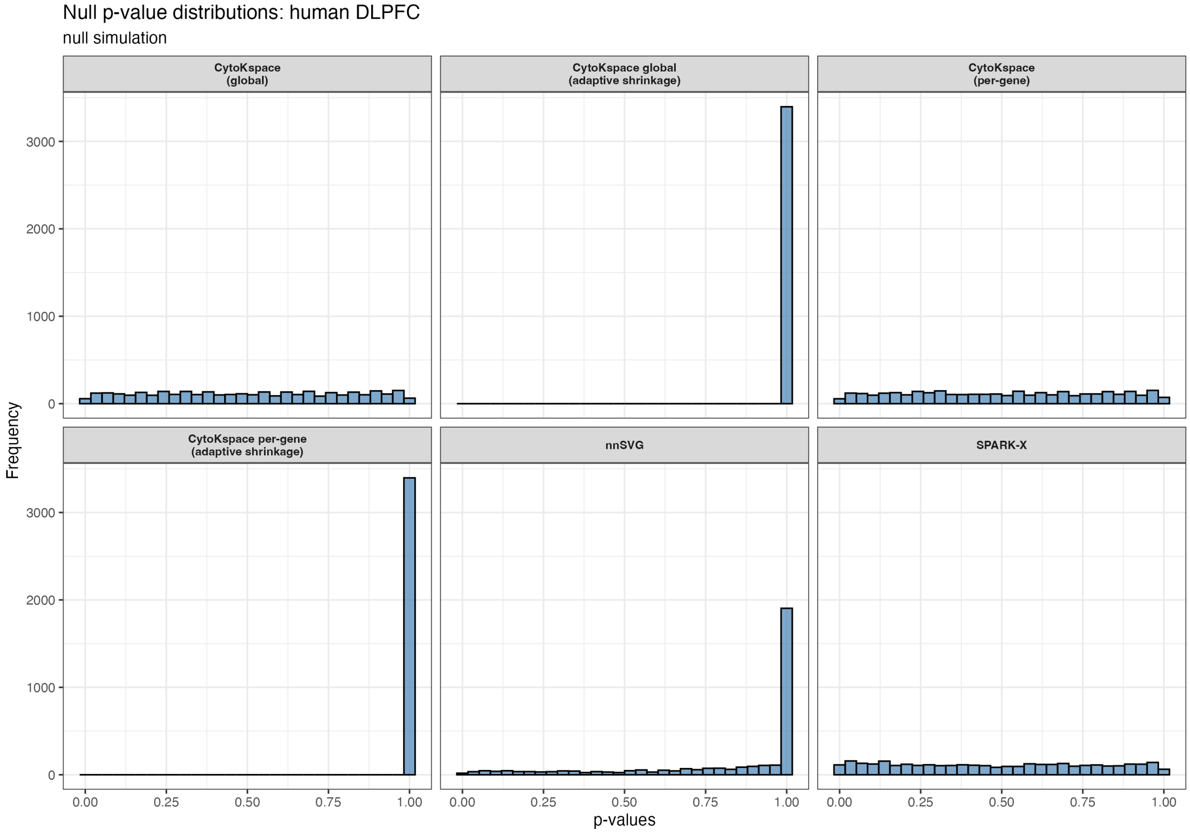


Figure S11: **Null $p$-value distributions on the LIBD human DLPFC dataset.** Six-panel facet of null $p$-value histograms from a real-data null simulation on the LIBD human dorsolateral prefrontal cortex Visium dataset (sample 151673; $N = 3{,}582$ spots, $G = 14{,}628$ genes after filtering). The null is constructed by independently permuting expression vectors across spots within the sample, breaking spatial signal while preserving the marginal expression distribution. Methods shown: CytoKspace (global), CytoKspace (global, adaptive shrinkage), CytoKspace (per-gene), CytoKspace (per-gene, adaptive shrinkage), nnSVG, and SPARK-X. CytoKspace (global/per-gene) and SPARK-X are well-calibrated (uniform on $[0, 1]$); nnSVG is mildly conservative; both adaptive-shrinkage variants concentrate mass near $p = 1$.

#### S2.12 Supplementary Figure S12: Null $p$-value distributions on the mouse OB dataset

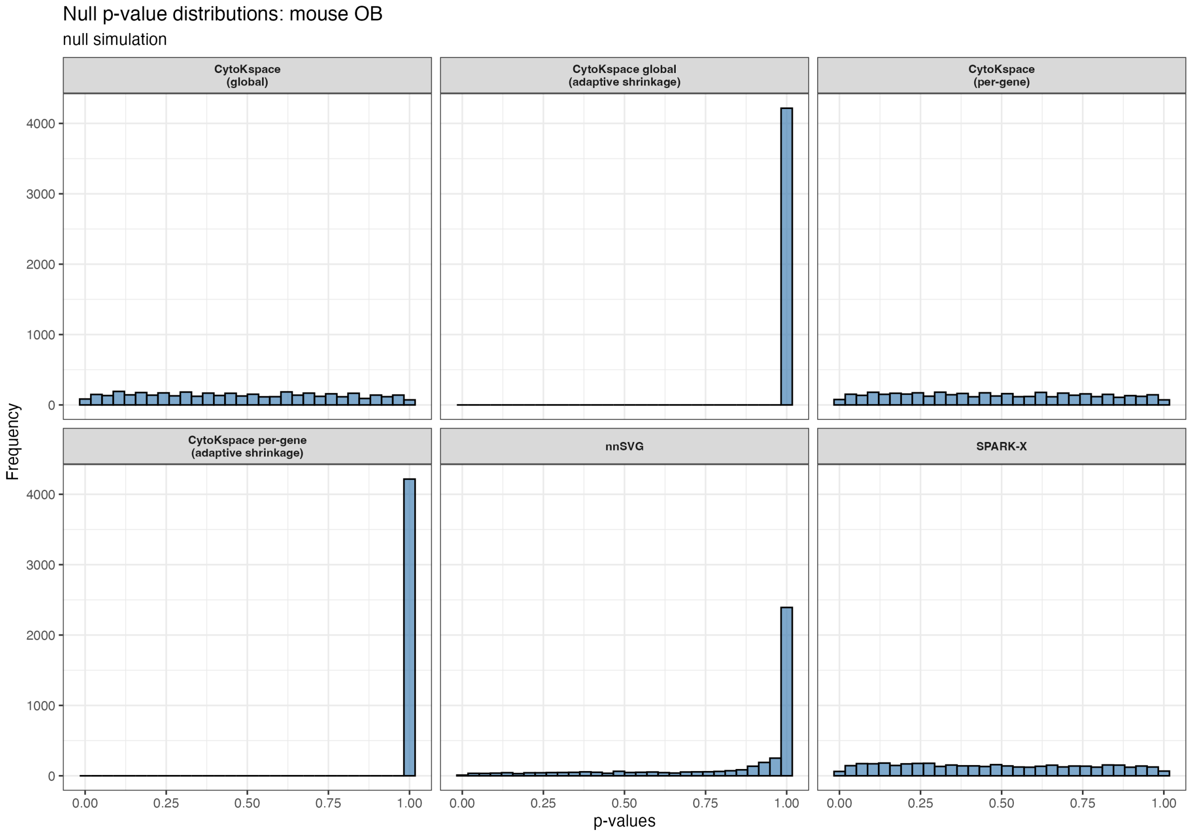


Figure S12: **Null $p$-value distributions on the seqFISH mouse olfactory bulb dataset.** Same display as Supplementary Figure S11 but on the seqFISH mouse olfactory bulb dataset ($N = 523$ spots, $G = 10{,}000$ genes) constructed via within-sample spot permutation. The qualitative pattern is identical to the LIBD case, demonstrating that CytoKspace's exchangeability-based inference remains well-calibrated on imaging-based platforms with very different noise characteristics from sequencing-based Visium (in particular, minimal zero inflation).

#### S2.13 Supplementary Figure S13: Adaptive shrinkage behaviour on the LIBD human DLPFC dataset

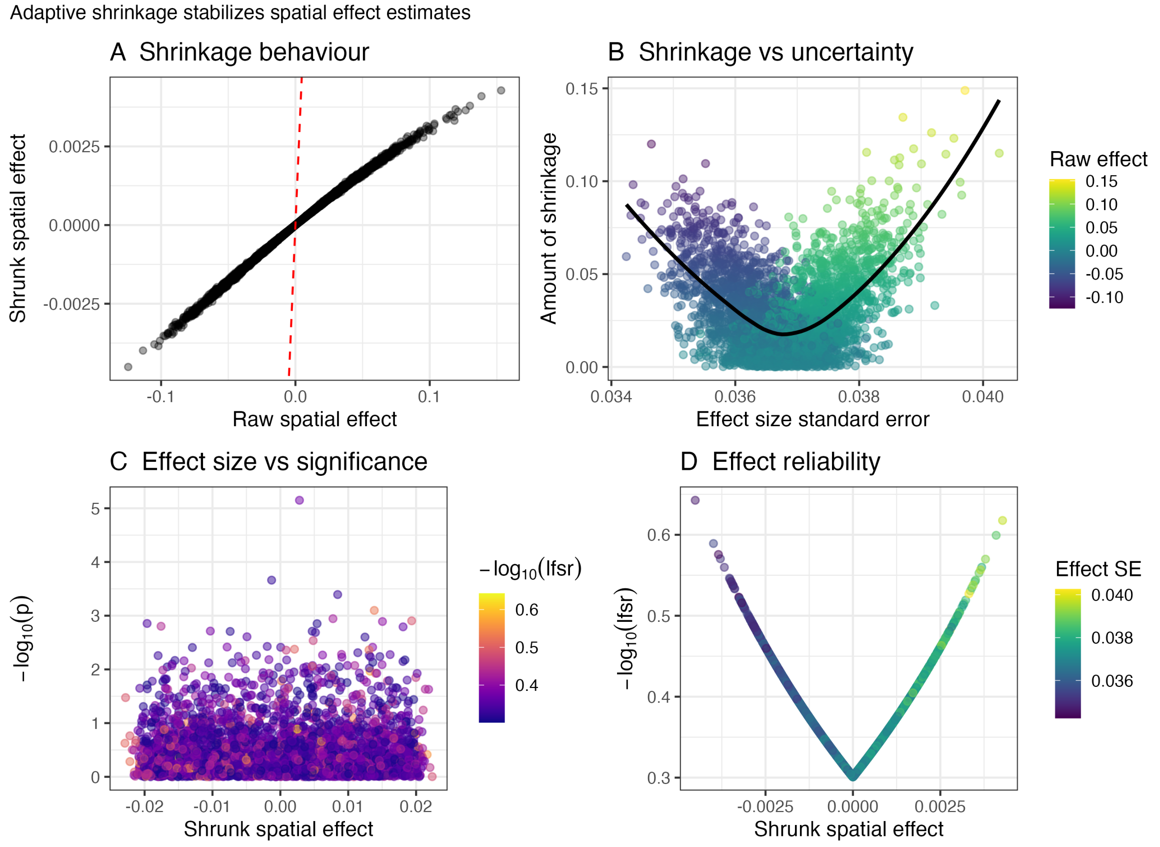


Figure S13: **Adaptive shrinkage stabilises spatial effect estimates on the LIBD human DLPFC dataset.** **(A)** Shrinkage behaviour: shrunk spatial effect $\hat{\theta}_g^{\text{ash}}$ versus raw spatial effect $\theta_g$ for all genes. The near-linear relationship with slope $< 1$ demonstrates systematic shrinkage toward zero, with the degree of compression increasing for genes with smaller raw effects. The red dashed line marks the origin. **(B)** Shrinkage versus uncertainty: amount of shrinkage ($|\theta_g - \hat{\theta}_g^{\text{ash}}|$) plotted against the effect-size standard error $\text{se}(\theta_g)$, with points coloured by the raw effect. The black loess curve confirms that genes with larger standard errors receive substantially more aggressive shrinkage, a mechanism that implicitly downweights the variance-inflated estimates identified by the mean-variance relationship (Shah et al. 2025; Law et al. 2014). **(C)** Effect size versus significance: $-\log_{10}(p)$ plotted against the shrunk spatial effect, coloured by $-\log_{10}(\text{lfsr})$. Genes in the upper tails combine high statistical significance ($p$-value) with high directional confidence (lfsr), whereas genes near the origin carry low significance regardless of their lfsr. **(D)** Effect reliability: $-\log_{10}(\text{lfsr})$ plotted against the shrunk spatial effect, coloured by effect-size standard error. The characteristic V-shape demonstrates that genes with larger absolute shrunk effects receive higher posterior directional confidence, with the lfsr penalty being most severe for genes near zero spatial effect (those most likely to be false positives or variance-inflated artefacts). Genes with small standard errors (dark) achieve high lfsr values at moderate shrunk effects, while genes with large standard errors (light) require substantially larger effects to achieve the same confidence, providing a principled gene-level correction for the mean-variance bias without requiring explicit observation-level precision weights.

#### S3 Extended benchmark tables

Table S1: Effect size sweep: full results, averaged across the $(N, G, \rho)$ simulation grid (main-text Section 3.1) and 100 Monte Carlo replicates per cell. Reference cell: $N = 1{,}500$, $G = 3{,}000$, $G_{\mathrm{SVG}} = 300$.

| $\delta$ | Method | Power | FDR | TP | FP | Runtime (s) |
|---|---|---|---|---|---|---|
| 0.3 | CytoKspace (global) | 0.800 | 0.050 | 240 | 13 | 42 |
| 0.3 | CytoKspace (global, adaptive shrinkage) | 0.680 | 0.010 | 204 | 2 | 42 |
| 0.3 | CytoKspace (per-gene) | 0.800 | 0.035 | 240 | 9 | 58 |
| 0.3 | CytoKspace (per-gene, adaptive shrinkage) | 0.740 | 0.015 | 222 | 3 | 58 |
| 0.3 | nnSVG | 0.800 | 0.030 | 240 | 7 | 2200 |
| 0.3 | SPARK-X | 0.760 | 0.035 | 228 | 8 | 2 |
| 0.6 | CytoKspace (global) | 0.970 | 0.058 | 291 | 18 | 44 |
| 0.6 | CytoKspace (global, adaptive shrinkage) | 0.880 | 0.008 | 264 | 2 | 44 |
| 0.6 | CytoKspace (per-gene) | 0.980 | 0.040 | 294 | 12 | 60 |
| 0.6 | CytoKspace (per-gene, adaptive shrinkage) | 0.960 | 0.007 | 288 | 2 | 60 |
| 0.6 | nnSVG | 0.970 | 0.010 | 291 | 3 | 2300 |
| 0.6 | SPARK-X | 0.810 | 0.042 | 243 | 11 | 2 |
| 1.0 | CytoKspace (global) | 1.000 | 0.045 | 300 | 14 | 46 |
| 1.0 | CytoKspace (global, adaptive shrinkage) | 1.000 | 0.005 | 300 | 2 | 46 |
| 1.0 | CytoKspace (per-gene) | 1.000 | 0.040 | 300 | 13 | 63 |
| 1.0 | CytoKspace (per-gene, adaptive shrinkage) | 1.000 | 0.003 | 300 | 1 | 63 |
| 1.0 | nnSVG | 1.000 | 0.018 | 300 | 6 | 2447 |
| 1.0 | SPARK-X | 0.830 | 0.045 | 249 | 12 | 2 |
| 1.6 | CytoKspace (global) | 1.000 | 0.042 | 300 | 13 | 46 |
| 1.6 | CytoKspace (global, adaptive shrinkage) | 1.000 | 0.002 | 300 | 1 | 46 |
| 1.6 | CytoKspace (per-gene) | 1.000 | 0.015 | 300 | 5 | 63 |
| 1.6 | CytoKspace (per-gene, adaptive shrinkage) | 1.000 | 0.001 | 300 | 0 | 63 |
| 1.6 | nnSVG | 1.000 | 0.015 | 300 | 5 | 2447 |
| 1.6 | SPARK-X | 0.830 | 0.078 | 249 | 21 | 5 |

Table S2: Multi-sample scaling: power and FDR across increasing $S$, averaged across the $(N, G, \rho)$ simulation grid (main-text Section 3.1) and 100 Monte Carlo replicates per cell. Reference cell: $N_s = 1{,}000$, $G = 2{,}000$, $G_{\text{SVG}} = 200$, $\delta = 0.8$.

| $S$ | Method | Power | FDR |
|---|---|---|---|
| 2 | CytoKspace Fisher (adaptive shrinkage) | 0.990 | 0.005 |
| 2 | DESpace (multi) | 0.850 | 0.065 |
| 3 | CytoKspace Fisher (adaptive shrinkage) | 0.990 | 0.000 |
| 3 | DESpace (multi) | 0.950 | 0.070 |
| 5 | CytoKspace Fisher (adaptive shrinkage) | 0.990 | 0.000 |
| 5 | DESpace (multi) | 1.000 | 0.075 |

Table S3: Comparison of multi-sample approaches: features, assumptions, and complexity.

| Feature | CytoKspace | DESpace | nnSVG |
|---|---|---|---|
| Sample adjustment | Within-sample residualization and standardization | NB model with sample intercepts $\gamma_{gj}$ | None (per-sample only) |
| Combination strategy | Fisher $p$-value combination + IV meta-analysis | Joint NB likelihood across pseudo-bulk counts | Average rank across samples |
| Inference type | Permutation (exact, finite-sample) | Parametric (edgeR quasi-likelihood, asymptotic) | Parametric (LR test, asymptotic) |
| Distributional assumption | None (exchangeability only) | Negative binomial | Gaussian |
| Spatial information | kNN kernel (continuous) | Pre-computed spatial clusters (discrete) | NNGP covariance (continuous) |
| Scalability ($S$ samples) | $O(GkN\bar{B}) + O(GS)$; embarrassingly parallel | $O(GN)$ joint fit; memory-intensive for large $S$ | $S\times$ single-sample; no combination |
| Cluster-level testing | No | Yes (individual cluster $p$-values) | No |
| Effect sizes | Shrunk $\hat{\theta}_g^{\text{ash}}$ + lfsr + posterior SD | Log fold-change per cluster | Proportion of spatial variance $\sigma^2/(\sigma^2+\tau^2)$ |
| Mean–variance correction | Not built-in (compatible with spoon weights) | Not addressed | Addressed by spoon (Shah et al. 2025) |

#### S4 Software and reproducibility

CytoKspace is implemented as an R package integrating with the Bioconductor `SpatialExperiment` class. The complete simulation framework is provided in `CytoKspace_simulations_v3.R`, which implements all benchmarks described in the main manuscript. Key implementation details:

- **Multi-sample permutations:** Within-sample permutations are implemented via serial `for` loops (not `bplapply`) to avoid closure serialization failures with `SnowParam` workers in BiocParallel.
- **Per-sample standardization:** Each sample is independently mean-centered and standardized to unit variance before computing the pooled $Q$ statistic, ensuring equal sample contributions regardless of expression magnitude.
- **DESpace comparison:** DESpace is called via `svg_test(spe_combined, cluster_col, sample_col, replicates = TRUE)` after combining samples with `cbind()`, which properly manages the `sample_id` slot. BayesSpace clustering is used when available; k-means fallback is provided.
- **Gene-order alignment:** Multi-sample wrapper functions explicitly reorder returned $p$-values to `gene_1, gene_2, ...` order via `match()`, ensuring correct alignment with the truth vector in `compute_metrics()`.
- **Dense count matrices:** Simulated counts are stored as dense `matrix` (not `dgCMatrix`) to ensure compatibility with DESpace/edgeR.
- **Unique column names:** Each sample's spots receive unique column names (`s1_spot_1`, `s2_spot_1`, etc.) so that `cbind()` for DESpace sees distinct samples.
